# Automated filtering of particle images in single particle cryoEM

**DOI:** 10.1101/2025.11.12.688030

**Authors:** Sony Malhotra, Daniel Hatton, Samuel Jackson, Matthew Iadanza, Agnel Praveen Joseph, Colin M. Palmer, Jeyan Thiyagalingam, Tom Burnley, Yuriy Chaban

## Abstract

Continued exponential growth in the number of structures resolved by single particle cryoEM, as seen in the last decade, requires ever more effective data analysis workflows. Datasets are rarely homogeneous, demanding a multistep procedure for discarding outliers. Since individual particles are very noisy, either 2D or 3D averages are normally used for discrimination. This becomes challenging when the 2D classes themselves are heterogeneous, leading to selection of contaminants or discarding useful rare views/poses. The 3D model-based discrimination requires trustworthy 3D maps and a correct assignment of Euler angles, which in turn depends on the quality of the initial data and might not be available at the very early stages of the analysis. We propose a novel deep-learning approach for improving quality of single particle datasets. The two-stage procedure consists of denoising single particle images using Variational AutoEncoder framework followed by particle quality filtering based on the score inferred for every particle by Domain Adaptation Neural Network trained on a large data set of categorised 2D averages. This approach allows an automated scoring of noisy raw images using data patterns learned from the high signal-to-noise ratio, externally derived 2D classes. Consequently, a higher quality data set enters computationally expensive steps of the data analysis, reducing the need for protracted and expensive calculations. Importantly, our method does not require any prior knowledge about the data or existence of a 3D model, making it universally applicable. Tests on publicly available datasets demonstrated that our approach largely outperformed 2D class-based particle discrimination. Smaller subsets of the top-scoring particles selected with our method were required to obtain the author-reported 3D model resolution. When applied to the user data in the automated on-the-fly data processing pipeline, the method rescued 30% of cases, which otherwise would not reach confidence threshold required for making decision to proceed to the 3D model refinement. It also led to general improvements in the quality of the 3D models for many datasets which were selected for the high-resolution processing.

## Introduction

Over the past decade, electron microscopy of cryogen-fixed, vitreous ice embedded specimens (cryoEM^1^) has been transformed by significant developments in both hardware and software. Most notable improvements are (semi-)automated sample preparation and handling, development of direct electron detectors with fast readout and shift to widespread usage of GPU (Graphical Processing Unit) in data analysis. Prominent software advancements include introduction of fast data collection schemes, motion correction, widespread adoption of maximum likelihood Bayesian frameworks and stochastic gradient descent-based methods for data analysis^2^. Taken together, these developments had synergetic effect and triggered the ‘resolution revolution’^3^, enabling the determination of atomic or near-atomic resolution structures of complexes at an exponential rate. CryoEM is currently the second most widely used method for structure determination (https://www.rcsb.org/stats/all-released-structures) and allows the study of biomolecules not amenable to x-ray crystallography or nuclear magnetic resonance spectroscopy. The Electron Microscopy database (EMDB, contains processed 3D cryoEM volumes and deposition is mandatory for most publications) has 49.5k entries and the Electron Microscopy Public Image Archive (EMPIAR^4^, contains raw data and deposition is optional) has 2423 entries (as of 18^th^ Sep 2025), and further rapid growth of both is widely anticipated.

To obtain a 3D map from an array of 2D snapshots, cryoEM images are processed using various software, including but not limited to RELION^5^, cryoSPARC^6^, EMAN2^7^, SPHIRE^8^ and CiSTEM^6^. Typical cryoEM image processing pipelines consists of the following steps: motion correction, contrast transfer function (CTF) estimation, particle selection and extraction, 2D classification and class selection, initial low-resolution 3D map generation, 3D classification, Euler angle refinement, and 3D model building and validation^9^. To efficiently use microscope time and handle the high volume of data collected using cryoEM, there is a need for automated image processing workflows, which can enhance the throughput and inform timely decisions by a microscopist. While initial steps of the data analysis are relatively quick and streamlined, the 3D analysis requires a multidimensional parameter search and usually takes much longer. Moreover, the quality of the data entering the 3D analysis determines its complexity and success. The biggest issues are caused by data set outliers, presence of poor-quality particles affected by air-water interface interaction artefacts, and preferred orientation. From the point of view of the Single Particle Analysis (SPA) workflow, the key contributors to the quality of the datasets are particle picking and the subsequent data cleaning routine.

Particle picking in cryoEM micrographs (images) can be done using several approaches. Usually, a combination of automated picking methods and manual interventions are used to optimize the resulting dataset. Peculiarities of a specimen, like uneven particle distribution, presence of aggregates and contaminants, together with typically low contrast and signal-to-noise ratio (SNR) of a cryoEM micrograph often complicate the task leading to inclusion of data outliers and underrepresentation of certain views^10^. To make informed decisions about cryoEM data quality, an increase in the SNR is generally achieved by averaging multiple images with similar characteristics^11^. In SPA, multiple aligned particles are used to compute 2D averages, which can provide a preliminary quality estimate by revealing potential preferred orientations of imaged macromolecules and the extent of data heterogeneity^12^. It is often acceptable to initially pick a larger number of particles, including ‘junk’ particles (*i.e.* those that would not contribute constructively to the final 3D reconstruction), and then perform further filtering in later steps of the pipeline, based on 2D or 3D class appearances, resolution and other metrics.

2D averaging is typically performed with a limited number of classes containing between a few tens to thousands of particles per class. Hence, the resulting 2D classes are usually heterogeneous. Prior to further processing, groups of perceived ‘junk’ 2D classes are removed during the class selection step, either manually or with automated class selection methods such as Cinderella^13^ (from the SPHIRE workflow^14^) or RELION class ranker^5^. However, both methods only classify the 2D class averages. In the case of heterogeneous 2D classes, filtering (or selecting) particles by removing (or retaining) whole class(es) is not necessarily reliable or efficient. An alternative method, CryoSieve, sorts individual particles based on the comparison of the high-frequency components of synthetic and observed particle images, and is reported to achieve high resolution maps with a selected subset of high scoring particles^15^. However, this method relies on prior knowledge of the 3D model and assumes that the particle Euler angles are correctly assigned, which is rarely the case during the initial stages of data analysis.

In this article, we present an automated deep learning-based particle filtering workflow comprising two steps, denoising and categorisation. The philosophy of our approach is to score and select individual particles following only an initial, reference-free 2D pose assignment, which is quicker and does not require a 3D model. Despite applying 2D poses, we utilise data complexity reduction for denoising of individual particles independently from 2D classification algorithms.

The denoising step uses an adaptation of a VAE (Variational AutoEncoder) framework, termed cryoVAE. Particles are then scored using classifier, cryoDANN (cryoEM Domain Adaptation Neural Network), which utilises knowledge from another domain, *i.e.* annotated 2D class averages, but applies this to the individual particle images in order to classify them. Domain adaptation is particularly useful in this context as it enables us to train the classifier on an annotated set from a different domain, in this case the higher SNR 2D classes, and then use it to make predictions for the denoised particle images. As a result, each particle image is assigned a quality score, which can be used to select particles that exceed a certain threshold, with this threshold potentially varying for different datasets. We benchmarked our approach by comparing quality of the datasets selected using cryoDANN, RELION class ranker^5^, manual 2D class selection by an expert, and randomly selected subsets of particles. We have tested the performance of cryoDANN using publicly available EMPIAR datasets, and user datasets collected and processed at the electron Bio-Imaging Center (eBIC)^16^ at Diamond Light Source. We demonstrate its usefulness in the data processing pipeline at eBIC and show how it can be used to filter out ‘junk’ particles and improve subsequent reconstructions for a variety of user datasets. Based on our findings, we propose that cryoDANN could serve as an additional automated particle sorting technique prior to *de-novo* 3D reconstruction and/or initial 3D classification to remove dataset outliers after the particle picking step.

The software is available at: https://gitlab.com/ccpem/cryodann under an open-source licence. We have also implemented the software for the cryoEM community through the CCP-EM Doppio software suite (https://www.ccpem.ac.uk/software), and it is integrated into the on-the-fly single particle analysis (SPA) data processing pipeline at eBIC, Diamond Light Source.

## Results

In this section, we describe our two-step automated particle filtering workflow. The first step is the denoising of the particle images. The second step uses a deep learning-based classifier which predicts the quality of the denoised images obtained from step 1. We then demonstrate the application and testing of the method on publicly available datasets from EMPIAR (https://www.ebi.ac.uk/empiar/) and how it compares to the other methods of data selection. We discuss the filtering of the author-deposited stack of particles (from EMPIAR) to achieve author-reported (to hundredths of an angstrom) resolution. Lastly, we showcase the practical considerations and benefits of the method for the early data processing stages based on the implementation of our method in a fully automated SPA data processing pipeline at eBIC, where it was run alongside standard RELION class-ranker approach.

### Particle filtering workflow and test dataset

The graphical overview of the workflow is shown in Figure 1a. The following is the order of steps applied on the set of picked particle images:

**Figure 1:**
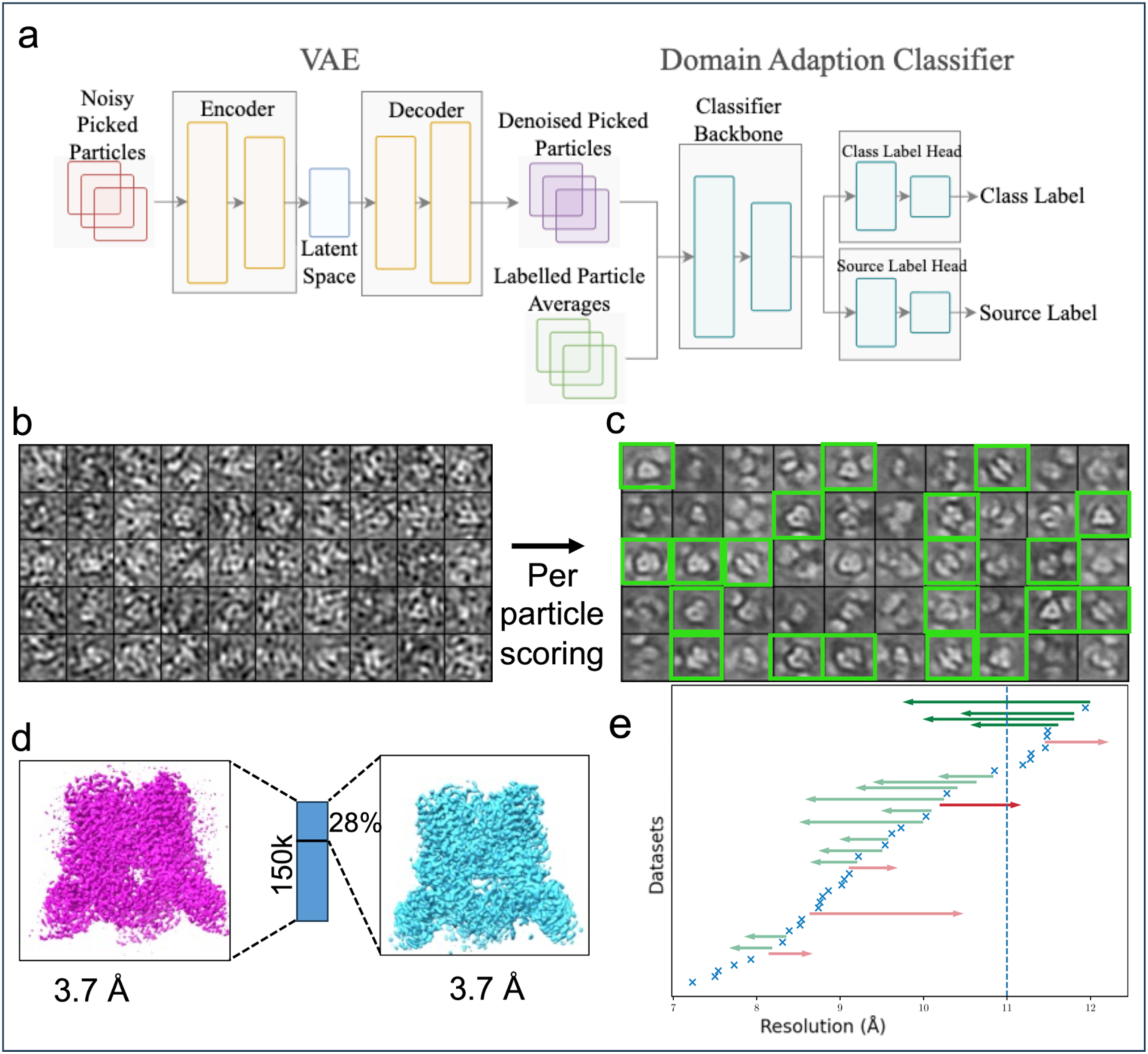
The overall workflow for denoising and scoring the picked particle images and its application. a. The model consists of an encoder-decoder architecture with a continuous latent space which can be used for reconstruction purposes. During the encoding part of the training process, each particle image is encoded into the low-dimensional latent space and then reconstructed using the learned latent variables. This results in denoising of the particle images which can be further used for feature extraction. The classifier is plugged in to learn these extracted features and assign a quality score to each particle image. b. The raw images used as an input to the c. The denoised particle images obtained after cryoVAE. The green squares show the particle images which were selected for further processing using cryoDANN scores. d. 3D reconstructions at the same resolution obtained using the whole set (pink) and ∼30% of the filtered particles (cyan). e. Change in 3D class resolution after standard RELION class-ranker score is combined with cryoDANN score in the automated on-the-fly data analysis pipeline. Resolution obtained after using RELION class-ranker particle filter is the starting point of an arrow; resolution obtained after application of the combination filter calculated as product of cryoDANN and RELION class-ranker score is the end point. Dotted vertical line shows 11 Å eligibility threshold for proceeding to a high-resolution 3D refinement. Blue crosses indicate data sets where both approaches resulted in the same resolution. Light-green arrows indicate data sets where the resolution improved within the 3D refinement selection boundaries; pink arrows indicate resolution decrease; blood-red arrow shows a data set where resolution decreased beyond the selection threshold; dark green arrows indicate resolution increase, which made a data set eligible for the high-resolution 3D refinement.

#### 1. Denoising the particle images

Based on reference-free 2D classification in RELION, in-plane transformations by real space interpolation were applied to the particle images, which were saved into particle stacks. These saved aligned particle stacks were used as an input to our Variational Autoencoder (cryoVAE) to obtain denoised particle images. CryoVAE is tailored to reconstruct the denoised particle stacks and generate latent vectors, which facilitates the visualization of particles within the learned latent space.

CryoVAE simultaneously learns an encoder which calculates a conditional distribution of a latent variable for a given stack of particle images, and a decoder which reconstructs a denoised image given this distribution of latent space variables. During the learning process, the latent space is regularized so that new images can be reconstructed using the decoder. The encoder tries to reduce the number of features for a given set of input images and compresses the data into encoded or latent space where data can be represented by latent space vectors (see Methods).

Following training, the output of cryoVAE comprises reconstructed (denoised) particle images (Figure 1b), learned latent vectors and a UMAP^17^ (Uniform Manifold Approximation and Projection) embedding of the latent vectors (a 2D histogram and a scatter plot, Figure S1).

#### 2. Using denoised images to classify particle images

We use the denoised particle images (obtained from cryoVAE, Figure 1b) as an input to a classifier: cryoEM Domain Adaptation Neural Network (cryoDANN). cryoDANN uses the knowledge from a different domain, *i.e.* the set of curated 2D classes, to assess the quality of the denoised particle images and outputs a score for each particle image. The network aims to minimize the classification loss for labelled source domain (the data on which it is trained *i.e.* 2D class averages) and domain confusion loss for all samples (see Methods for details).

The two different losses/accuracies are:

- Classification loss/accuracy: this is the loss/accuracy of the class label head on the labelled dataset (2D class averages in this case, see Methods for more details)
- Domain loss/accuracy: this the loss/accuracy of the source label head in distinguishing between the two domains (Domain 1: labelled 2D class averages, Domain 2: unlabeled images from cryoVAE).

The computed score can then be used to filter out low-quality particles (Figure 1c) which would not contribute constructively to a 3D reconstruction.

We tested the entire workflow (as shown in Figure 1a) using four EMPIAR datasets (Table 1), which were processed as described using the data processing pipeline described in Methods:

**Table 1:** The datasets used for testing the performance of our particle filtering approach.

| **Data set** | **EMPIAR entry** | **Number of micrographs** | **Voltage (keV)** | **Pixel size (Å)** | **EMDB ID** | **Resolution (Å)** |
| --- | --- | --- | --- | --- | --- | --- |
| Beta-galactosidase | 10204 | 1338 | 300 | 0.885 | 6840 | 2.6 |
| Jack bean Urease | 10547 | 239 | 200 | 0.83 | 20016 | 2.8 |
| Cannabinoid receptor | 10288 | 2756 | 300 | 0.86 | 0339 | 3.0 |
| GAPDH | 10955 | 270 | 300 | 0.813 | 32162 | 2.2 |

Both aligned (in plane transformations by real space interpolation, using the “--apply_transformation” flag for the “relion_stack_create” program) and unaligned particle sets were used to test the performance of the denoiser (cryoVAE). When the aligned set of particles was used as input, we observed distinct clusters in the latent space and the denoising was more efficient (Figure S2). Therefore, the initial alignment of the particle stack is crucial for the current state of cryoVAE to put all particles in register according to the assumed pose but does not use the 2D class averages themselves. We do not see any meaningful training and representation in latent space with the unaligned set of particles (latent space visualization, Figure S2). For the test dataset, the loss values (both class and source domain loss) along with class and source domain accuracies are plotted per epoch in Figure S3.

After applying our workflow (denoising followed by per particle scoring, Figure 1a), we demonstrate that processing the obtained filtered set of particles (∼28% in this case, Figure 1d) results in the same resolution reconstruction as with the full set of particles (150k, see section “Assessment of cryoDANN on EMPIAR datasets with particle stacks” for more details). This implies that using cryoDANN scores to discard a good proportion of ‘junk’ particles very early on during the data processing does not compromise the quality of the reconstruction obtained (Figure 1d). After testing the performance and versatility of our score on multiple datasets we implemented our data filtering methodin the automated data processing pipeline at eBIC. At eBIC, on-the-fly data processing is used to give users real time feedback on data quality and provide with starting point for advances data analysis. The start of a 3D refinement is conditional on one of four 3D classes calculated from a dataset of ∼200,000 coarsened particles reaching at least 11 Å resolution, hence the dataset should contain a sufficient number of good particles. In most scenarios, no assumptions are made about the 3D model, and de-novo 3D reconstruction is run for every data set using best 2D classes selected with RELION class-ranker. Our tests revealed that using cryoDANN score in combination with RELION class ranker score, but without any dataset specific optimisation performs better (better resolution for the best 3D class) than the existing class selection in the pipeline (Figure 1e, for more details see section “Assessment of cryoDANN enhanced pipelines”). Crucially, for 4 datasets the resolution improved sufficiently to warrant their high-resolution refinement (this is 30% of all data with the initial best 3D class resolution worse than 11 Å).

### Performance on the test dataset and comparison with other methods

To find an efficient cryoDANN score (classification probability) threshold for particle filtering, we arbitrarily split classes into ‘good’ and ‘bad’ categories based on their SNR, which often is reflective of the number of a particles in a 2D class. In Figure 2, we highlight this for one of the test datasets: EMPIAR-10204. The cryoDANN score distribution for all the particles in this batch is shown in Figure 2a. Particles are given a score between 0 and 1, with higher scores implying better particle quality. The number of particles per RELION 2D class is shown in Figure 2b. As most of the particles have a score less than 0.1 (Figure 2b), we assessed five score thresholds 0.05, 0.06, 0.07, 0.08 and 0.09. It should be noted that these cutoff values are very low because the training of cryoDANN is performed using the images of class averages, which have a higher signal to noise ratio than a single particle image. Also, as the 50k particle subsets are obtained using standard picking hence they contain many low scoring ‘junk’ particles (Figure 2b, score distribution skewed to the left).

**Figure 2:**
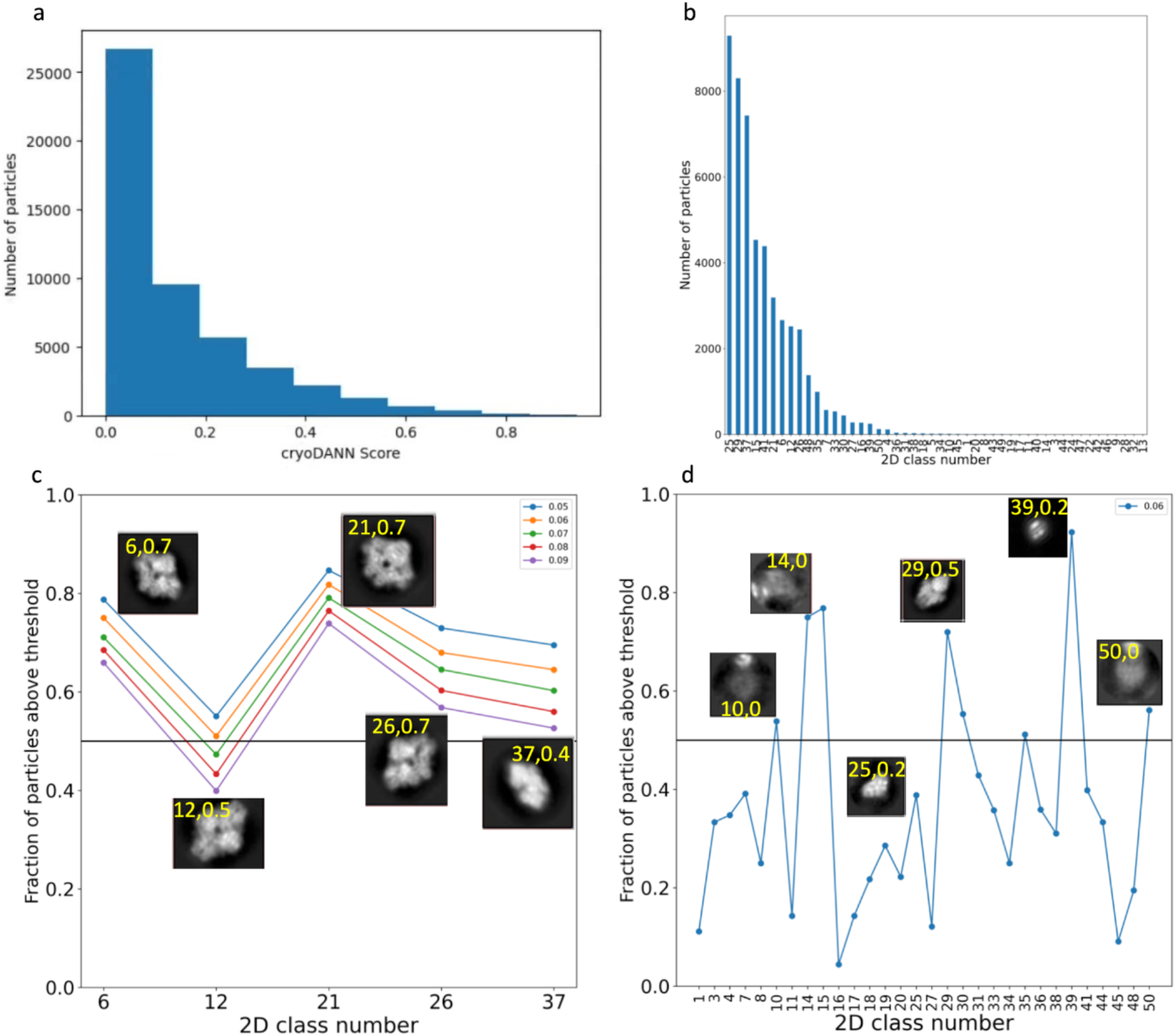
Particle filtering and score threshold for EMPIAR-10204. a. cryoDANN score distribution for a subset of 50k particles from the eBIC data processing pipeline. b. The distribution of particles in RELION 2D classes. c. and d. The proportion of particles in each class that exceed the cryoDANN score threshold, shown for manually selected ‘good’ classes and for the remaining classes categorized as ‘others’. The class number and RELION class ranker score are in yellow.

For each of the five manually selected ‘good’ classes, at least 50% of the particles scored greater than the selected threshold of 0.06 (Figure 2c). For the rest of the classes, 73% of the classes have less than 50% of the particles scored below the threshold of 0.06 (Figure 2d). The 2D classes in Figure 2d such as class number 29 and 21 appear heterogeneous in nature and hence are a mix of ‘good’ and ‘bad’ particle images. This suggests that a threshold of 0.06 provides a useful initial estimate of particle quality in this case. This threshold may vary for other datasets depending on the quality of particle images and training performance. In EMPIAR datasets 10024, 10288 and 10547, most particles from manually selected good classes score above the 0.06 threshold, while most from bad classes score below it. Interestingly, in the fourth dataset (EMPIAR-10955), the 11k particles selected at a threshold of 0.06 yield a resolution comparable to that obtained using the 26k particles from the manually selected ‘good’ classes (Table 2). However, most particles from the manually selected ‘good’ classes do not score above the threshold (Figure S4), which suggests that the ‘good’ classes may be heterogeneous, highlighting the need for objective particle scoring and selection methods such as cryoDANN.

**Table 2:** Number of selected particles used for reconstructions.

| EMPIAR-ID | Number of selected particles at selected threshold (0.06) | Number of particles in manually selected good classes |
| --- | --- | --- |
| 10204 | 29501 | 32189 |
| 10547 | 30702 | 34700 |
| 10288 | 23403 | 32189 |
| 10955 | 11136 | 26559 |

For each of the four datasets, we randomly selected a batch (one of the 50k particle subsets obtained from the eBIC data processing pipeline, see Methods) to compare the performance of cryoDANN with RELION class ranker, manually selected good classes, and a randomly selected subset of particles (Figure 3). To compare these metrics meaningfully, the reconstructions need to be performed with similar number of particles. Therefore, for a fair comparison, if cryoDANN selected N particles out of the 50k subset, we selected the top N scoring particles from class ranker and N randomly-selected particles. An expert-selected set of classes was used to obtain a manual selection of particles. Table 2 lists the number of particles selected at a cryoDANN threshold of 0.06 and in the manually selected good classes.

**Figure 3:**
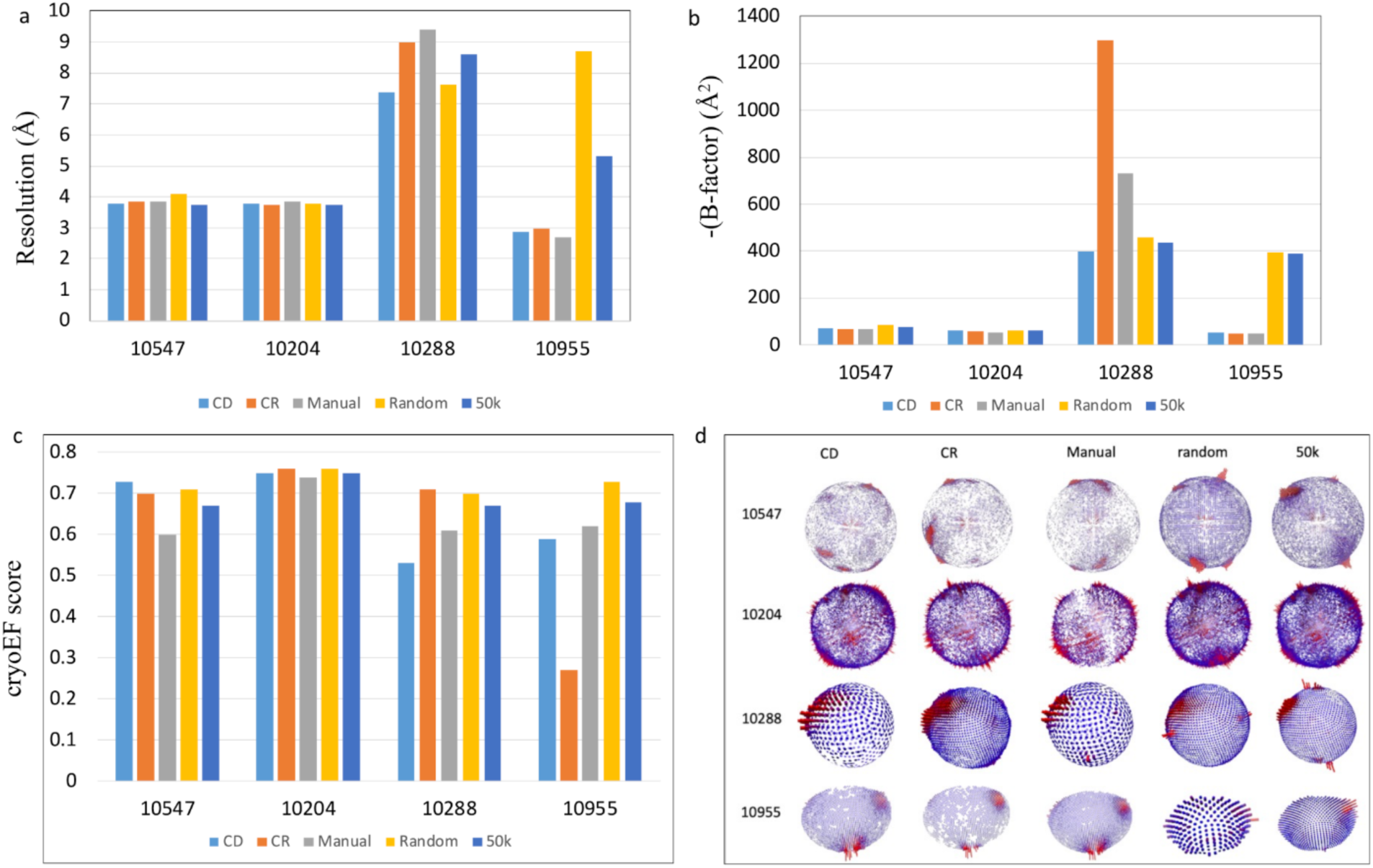
Performance of cryoDANN on four EMPIAR datasets processed using the eBIC data processing pipeline. The metrics (a) resolution, (b) B-factor (Rosenthal and Henderson, 2003) and (c) cryoEF score were used to assess the performance. d. Distribution of Euler angles for 3D reconstructions obtained using different subsets of particles. The larger red cylinders represent preferred orientations (orientations comprising more particles).

CryoDANN’s performance was better or comparable to the manually selected good classes, as judged by the resolution of 3D refinement (Figure 3a). The B-factor also suggests that the performance of cryoDANN filtered particles is comparable to manual filtering (selection of good 2D classes, Figure 3b). Comparisons with the full set of 50k particles suggest that automated filtering of particles using cryoDANN results in better resolution, lower B-factors and comparable distribution of Euler angles (Figure 3c and 3d for EMPIAR 10955) computed using cryoEF^18^.

Figure 4 shows reconstructions generated for EMPIAR-10547 using filtered particle sets from the different methods, namely: RELION class ranker; cryoVAE and cryoDANN; manual selection of classes; randomly selected particles. The reconstruction obtained using the full subset (50k) is also shown in Figure 4e for comparisons. Filtered particles obtained using cryoDANN (Figure 4b) result in a comparable quality reconstruction as manually selected good particles (Figure 4d).

**Figure 4:**
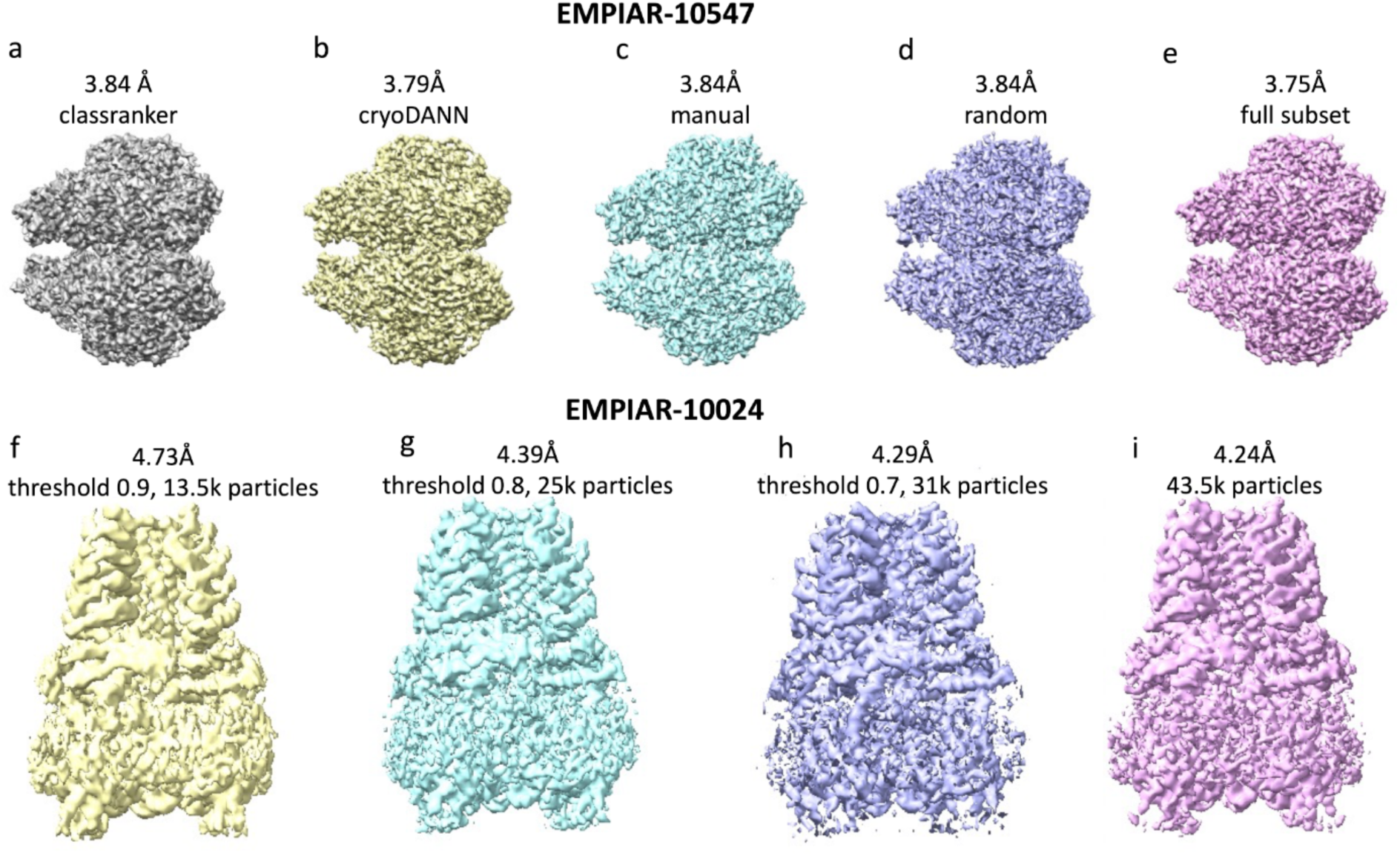
Reconstructions obtained using EMPIAR datasets. EMPIAR-10547-One of the subsets of 50k particles obtained for EMPIAR-10547 using eBIC data processing pipeline were filtered using different methods and reconstructions obtained and labelled as a. RELION class ranker, b. cryoDANN selected particles, c. manually selected good classes, d. randomly selected particles, and e. full subset of 50k particles. The bottom panel shows the particle filtering for EMPIAR-10024. f. Particle stack (with 43,585 particles) was downloaded and refined using RELION. Different score thresholds were tested for cryoDANN (0.9,0.8 and 0.7). g. Using the top 13.5k particles (at score threshold of 0.9), we could obtain a resolution of 4.73Å. h and i. As the score threshold was relaxed to 0.8 and 0.7, the number of particles increased to 25k and 31k respectively. However, the resolution obtained was 4.39 Å and 4.29 Å respectively.

For the dataset EMPIAR-10288, we could not obtain a reasonable resolution reconstruction from the 50k particle subset. Even with all of the 50k particles in the batch, the achieved resolution is only 8.6 Å (although filtering using cryoDANN still results in a better resolution of 7.4 Å). A larger particle set might be necessary to obtain a higher resolution reconstruction with difficult datasets like this.

### Assessment on EMPIAR datasets with particle stacks

The above datasets were all processed with a subset of 50k particles using the eBIC on-the-fly automated data processing pipeline. This pipeline uses the crYOLO^14^ particle picker for an automated particle selection, which also filters particles for “quality” and SNR based on a generally-trained model. We tested our method on the five publicly available datasets from EMPIAR^4^ which have author deposited particle stacks. The availability of particle stacks removes any subjectiveness of our particle selection and choices during the data processing. It also allows us to test the performance of our approach on pre-selected particles.

We subjected the author-deposited particle stacks to filtering using cryoDANN. We then performed two 3D refinement jobs in RELION, using all particles *i.e.* the author-deposited particle stack (Table 3, column 4) and a filtered set of particles obtained from cryoDANN (Table 3, column 6). For these EMPIAR datasets, we believe these particles have already undergone some level of manual filtering by the authors before deposition. We were able to filter these particles further and obtain a resolution similar (to within hundredths of an angstrom) to the value obtained from the whole stack of particles. For one of the cases, EMPIAR-10255, we were able to obtain the author-reported resolution by using only 28.4% of the particles, which clearly demonstrates the benefits of using our method for data set screening. Different thresholds tested for these datasets and the resolutions and B-factors obtained are provided in Table S1.

**Table 3:** EMPIAR datasets used for assessing the performance of cryoDANN in filtering of the particles.

| EMPIAR ID | Author-reported resolution (Å) | Number of particles | Resolution value calculated in-house with all particles(Å) | Filtered number of particles | Resolution with filtered particles (Å) | % of particles retained |
| --- | --- | --- | --- | --- | --- | --- |
| 10024 | 4.24 | 43585 | 4.24 | 31462 | 4.29 | 71.9 |
| 10097 | 4.24 | 130000 | 4.24 | 88099 | 4.29 | 67.8 |
| 10093 | 3.55 | 175344 | 3.74 | 155390 | 3.76 | 88.6 |
| 11233 | 3.2 | 42071 | 3.19 | 32761 | 3.19 | 77.8 |
| 10255 | 2.9 | 157224 | 3.7 | 44682 | 3.77 | 28.4 |

Figure 4f-i highlights this for one of the datasets, EMPIAR-10024. This is a 4.24 Å structure of TRPA1 channel with its agonist. We used the full deposited stack of particles (43.5k) as a positive control and performed in-house refinement followed by RELION post processing. We obtained a resolution of 4.24 Å, which is equal to the published resolution (Figure 4i). We denoised the particle stack using cryoVAE and performed filtering on the denoised stack using cryoDANN. Using the 13.5k top-scoring particles (score threshold 0.9), we obtained a resolution of 4.73 Å (Figure 4f). Adding more particles by relaxing the score threshold to 0.8 and 0.7, we obtained resolutions of 4.39 Å and 4.29 Å respectively (Figures 4g and 4h). Therefore, using our method the selected set of particles can further be pruned without a significant drop in resolution. To further assess the quality of filtered-out (low-scoring) particles, we performed 3D refinement jobs with the lowest-scoring 12k and 18k particles, corresponding to the thresholds of 0.7 and 0.8 respectively. This resulted in the reconstructions with resolution of 5.44 Å and 4.67 Å respectively, implying that low scoring particles are less informative than the high scoring particles (Table S1).

It is important to note that these particles stacks obtained from EMPIAR are the final particles which went into the reconstruction, hence they may have been carefully selected by the authors after iterative rounds of either 2D or 3D classification. This may lead to the following two observations:

1. The scores distribution for these (author curated) particle images are skewed towards the higher values as compared to those obtained from the 50k-particle batches (see section Performance on the Test Dataset). We believe this is because these are already filtered particles (used for final reconstruction by the authors) and hence are better quality particle images, as opposed to the 50k particles, which were obtained through a standard automated picking process.
2. The resolution and B-factors obtained illustrate the dependence on the number of particles rather than the quality of particles. This is further very clearly noted in Table S1. For EMPIAR-10024, the author-deposited stack of ∼43.5k particles results in a resolution of 4.24 Å. With a decreasing cryoDANN threshold (increasing number of particles), we see a slight improvement in the resolution obtained. An equal number of randomly selected particles results in comparable resolution, which implies that the author deposited particle stack is of very good quality, i.e. all the particles are ‘good’ and contribute approximately equally to the final reconstruction. Because of the quality of the final particle stack, we also experimented with removing the denoising step (cryoVAE) and only running the cryoDANN (classification) step on the original particle images. As expected, we note that the top-scoring particles filtered using cryoDANN result in a similar resolution as with the denoising step. This implies denoising significantly enhances classification performance for raw or noisy particle images, but may be optional when processing high-quality, pre-filtered particle stacks (Table S1). The same trend is also observed for EMPIAR-10097 where a randomly selected subset of ∼60% of the particles resulted in a similar resolution as obtained with the full set of particles (Table S2).

For EMPIAR-10093, we observe that over 80% of the particles must be retained to obtain a resolution similar to the full set of particles, implying that the particles are not dispensable in this case. To investigate this, we compared the highest-scoring particles with the lowest-scoring ones. For instance, the top 84k particles achieve a reconstruction at 4.08 Å, while the lowest scoring 90k particles yield a reconstruction at 4.38 Å (Table S3). EMPIAR-11233 has a particle dataset with only 42k particles of high quality, where we observed a retention percentage of 77.8% was needed to achieve the author-reported resolution (Table S4). Therefore, for this dataset we did not perform a comparison with lowest scoring particles.

For EMPIAR-10255 there are two particle stacks deposited in EMPIAR: all the picked particles (900k) and the 157k particles which were used to build the final map (EMD-0593). The reported resolution of this structure (EMD-0593) is 2.9 Å using the 157k particles. However, using eBIC’s automated data processing pipeline and the mask built using EMD-0593 for RELION Refine3D, we obtained a resolution of 3.7 Å. We used this resolution value to compare the performance of our particle filtering approach. By using only the top 28% of the particles from this set of 157k particles, we obtain a similar resolution, hence demonstrating the application of our filtering approach (Tables 3 and S5). However, in the case of this data set, similar resolution can be obtained by random selection of the same number of particles. This indicates that the deposited set of particles is of particularly high quality and was likely carefully selected by the authors.

Therefore, to further assess our filtering of particles, we used the set of all picked particles (900k) for EMPIAR-10255 (Table S6). Using our filtering method, we selected the 66k top-scoring particles (∼7% of the total particles) and were able to obtain a resolution of 3.7Å. This is the resolution we obtain using the particles that the authors used for their final refinement (157k particles, ∼17% of total data). It is important to note that the filtered-out particles may exhibit different conformation(s) of the molecule, implying heterogeneity.

### Data analysis pipeline efficiency enhancement using cryoDANN

One of the aims of particle filtering methods, such as the one described above, is to enhance automated processing pipelines, providing unsupervised removal of poor quality or misidentified particles that negatively impact reconstruction quality. In order to assess what improvements can be gained from the incorporation of cryoVAE+cryoDANN into automated processing we investigated its performance on the processing pipeline deployed at eBIC (discussed in more detail in the Methods section).

We are interested here in the estimation of the performance of different particle filtering techniques while keeping the rest of the pipeline the same. We therefore distinguish the pipelines based on the particle filter, *f*, that they use. The different filters that we tested on 50 user datasets (for selection criteria see Methods) are:

1. Particle filter (*f_a_*), is the original pipeline which uses the results of the RELION class-ranker to reject aprox. 50% of the particles. If a particle is a “member” of a 2D class that is scored below a threshold (determined from the analysis of the first batch comprising of 50,000 particles) then it is excluded from further analysis.
2. Filter (*f_b_*), in which cryoVAE is trained on each batch (subset of 50k particles) and then each denoised particle image is scored by cryoDANN. (Note that 2D classification is still required in this workflow to obtain estimates of the particle alignments relative to each other in a given class.). As in (1.), aprox. 50% of particles are rejected.
3. Filter (*f_ab_*), which is a combination of the two filters (f*_a_* and f*_b_*) described above. Each particle is scored as the product of the scores obtained from class ranker and from cryoVAE+cryoDANN. The top 50% of particles are selected based on that combined score.

The performance for these filters is summarised in Figures 5 and S5. When comparing the class ranker-based pipeline (using filter *f_a_*) with the naïve implementation of a cryoVAE+cryoDANN pipeline (filter *f_b_*), approximately one third (17) of the datasets, referred to as subset *f_a_* = *f_b_* (filter *f_a_* performance is equal to that of filter *f_b_*), showed equivalent performance according to the metric of best 3D class resolution. In these cases, there is on average a sub-10% shift in cryoEF^18^ efficiency score (Figure 5b). 10 of these 17 see an improvement in cryoEF score for *f_b_*. In 42% of cases (21 datasets), referred to as subset *f_a_* > *f_b_*, the existing class ranker-based pipeline outperformed the naïve cryoVAE+cryoDANN pipeline. However, in 16/21 cases the cryoVAE+cryoDANN pipeline leads to an improvement in cryoEF efficiency score. For the remaining 12 cases the cryoVAE+cryoDANN pipeline outperforms the class ranker pipeline in 3D class resolution, referred to as subset *f_b_* > *f_a_*. 9 of these also have improved efficiency scores.

**Figure 5:**
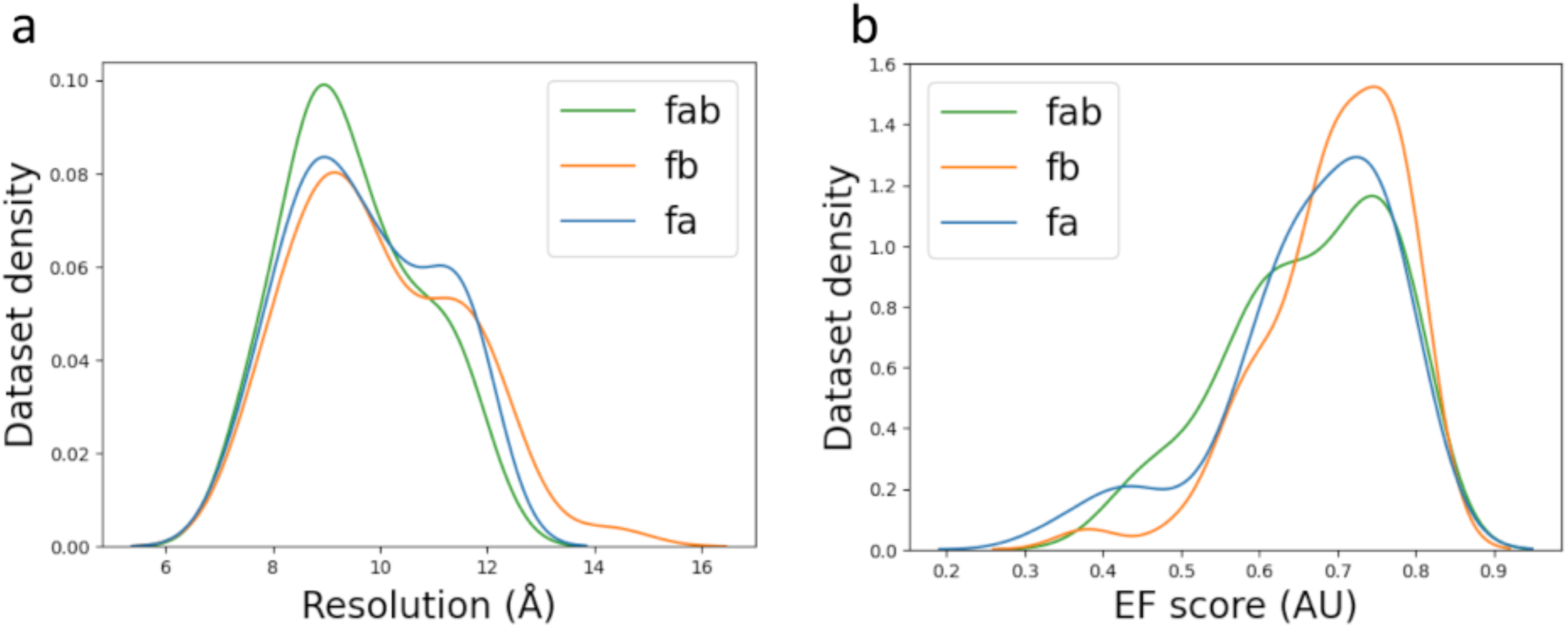
Performance in the automated data processing pipeline. The existing particle filter *f_a_* (blue, using the RELION class ranker) is compared to filter *f_b_* (orange, using cryoVAE and cryoDANN) and the combination filter *f_ab_* (green, using product of *f_a_* and *f_b_* scores). The distribution (kernel density estimation) shows the a. resolution and b. cryoEF score of the best 3D class for the 50 user datasets collected at eBIC.

The subset of datasets where filter *f_ab_* achieves the same best 3D class resolution as *f_a_* will be referred to as *f_a_* = *f_ab_*, the subset where *f_a_* outperforms *f_ab_* will be referred to as *f_a_* > *f_ab_* and the compliment subset where *f_ab_* outperforms *f_a_* will be referred to as *f_ab_* > *f_a_*. Subset *f_a_* = *f_ab_* is larger than *f_a_* = *f_b_* at 60% of the total (Table 4). The increase here largely comes from datasets that are in *f_a_* > *f_b_* indicating an improvement of *f_ab_* over *f_b_* for these cases (Figure 5a). There are 4 datasets in both *f_a_* > *f_b_* and *f_ab_* > *f_a_* indicating the ability of *f_ab_* to improve on cases where the class ranker is performing well on its own. There are 3 datasets which are in both *f_a_* = *f_b_* and *f_ab_* < *f_a_,* showing that there are cases where the hybrid approach makes an improvement on data which performs equally well under cryoVAE+cryoDANN and the class ranker on their own.

**Table 4:** Performance comparison of automated data processing after implementation of cryoDANN. The subsets are defined, and their sizes are listed in this table for easier interpretability.

| Subsets | Description | Size |
| --- | --- | --- |
| $f_a = f_b$ | Similar performance when comparing filters $f_a$ and $f_b$ | 17 |
| $f_a > f_b$ | $f_a$ outperforms $f_b$ | 21 |
| $f_b > f_a$ | $f_b$ outperforms $f_a$ | 12 |
| $f_a = f_{ab}$ | Similar performance when comparing $f_a$ and $f_{ab}$ | 30 |
| $f_a > f_{ab}$ | $f_a$ outperforms $f_{ab}$ | 5 |
| $f_{ab} > f_a$ | $f_{ab}$ outperforms $f_a$ | 15 |

The average resolution difference between the two pipelines in subset *f_a_* > *f_b_* is 1 Å. For subset *f_b_* > *f_a_* this is 0.74 Å. This assessment shows that there are cases in which filter *f_a_* outperforms filter *f_b_* and vice versa, but overall filter *f_a_* performs better according to these metrics. This motivates the use of filter *f_ab_* described above. This combined filter will balance the two approaches (as seen from the distribution of resolution of best 3D class and the cryoEF score in Figure 5) by removing low quality particles from classes which score high in class ranker but allowing the rescue of high-scoring particles from the classes which would be otherwise rejected by class ranker.

A summary of the comparison results for *f_b_* and *f_ab_* is shown in Figure S5. Here the best 3D class resolution and the cryoEF efficiency score for *f_b_* and *f_ab_* are divided by the value obtained from *f_a_*. The resolution obtained from a filter *f_i_* is denoted *R*(*f_i_*) and the efficiency score is denoted *E*(*f_i_*). The ratio is inverted for resolution so that a larger value indicates an improvement over the baseline case of *f_a_* as is the case for the efficiency score. The tabulated results for the resolution across the three different pipelines are given in Table S7.

For progression to the 3D refinement stage in the eBIC data processing pipeline, a 3D classification resolution threshold of better than 11 Å is required. The combined approach (*f_ab_*) was able to recover 30% of datasets (15% with *f_b_* alone) that had previously failed to advance due to resolution limitations in the automated pipeline (Table S7).

In terms of computational time for each batch (50k particles), on average cryoVAE training took 8 minutes on a single GPU (mixture of P100s and V100s) with cryoDANN inference taking 3 minutes on average.

## Discussion

In this article, we have presented an automated deep learning-based particle filtering workflow. This workflow begins with denoising followed by classification and assigns a quality score to particle images. We have tested our method on publicly available datasets from EMPIAR and demonstrated that particles can be successfully filtered using this approach, which can then be used to perform 3D refinement. Our approach can be used while processing the data and selecting the ‘good’ particles for downstream data processing steps. This overcomes the 2D class heterogeneity problem as the particles can be individually scored as opposed to the class-based automated selection/sorting enabled by RELION class ranker.

It is up to the user’s discretion to decide how many particles they wish to retain and the score threshold will vary depending on the quality of the dataset. Based on our observations, selecting the top 50% of particles ranked by model confidence consistently yields good performance when evaluated within an automated data processing pipeline. It is fair to assume that this suggestion is less likely to get rid of underrepresented poses/conformations, the fate of which should be decided at later stage of the data analysis. This seems to be especially important for the cases where most picked particles are good already.

We have made the software freely available for academic use, and a user interface is implemented in the CCP-EM Doppio software package (Figure S6).

In summary, our approach provides a method to discriminate good particle images from junk images during the initial stages of the data processing pipeline and therefore makes the downstream steps more efficient and faster.

We have performed an initial implementation of an automated processing pipeline using cryoVAE+cryoDANN particle filtering at the UK national cryoEM facility, eBIC, at Diamond Light Source, and compared this to a current pipeline which uses only the RELION class ranker for particle filtering. The cryoVAE+cryoDANN particle score for filtering in the eBIC SPA pipeline (which was optimised with the RELION class ranker in mind) produced mixed results. However, the combination of cryoVAE+cryoDANN and RELION class ranker scores (combined pipeline) leads to more cases where an improvement is made over the class ranker on its own and significantly fewer cases where the class ranker pipeline performs better. By resolution metrics, the combined pipeline performs the best of the three on average across 50 datasets. There are also notable cases where the cryoVAE+cryoDANN pipeline, and the combined pipeline, produce significant improvements in the distribution of angular assignments of particles. Furthermore, in 30% of the datasets, the combined pipeline was able to recover cases that previously failed in the existing workflow (best 3D class resolution worse than 11 Å), enabling progression to 3D refinement.

CryoVAE+cryoDANN is now implemented at eBIC to improve the efficiency of users’ data collection sessions and is suitable for deployment elsewhere at similar high throughput cryoEM centres. With this approach, we will be able to provide improved feedback to the users on the quality of data while data collection is in progress. Additionally, we will continue to actively develop this method, using the results from testing on additional datasets to further refine and tune our deep learning model.

Here, we described a novel approach for streamlining single particle selection using Domain Adaptation. This allows an automated categorisation of noisy raw images using data patterns learned from high signal-to-noise ratio, externally derived 2D classes. It is interesting to note, that the method reflects on decades of user experience pre-dating fast data acquisition techniques, where relatively small data sets were interactively categorised to obtain the best result from limited resources. We hope that our approach, which utilises deep-learning techniques can now bring the same high standards into the era of high throughput cryoEM data processing.

## Methods

Our method uses a two-stage machine learning (ML) pipeline for processing picked cryoEM particles. Importantly, our method makes decisions at the individual particle image level, in contrast to other available methods which categorise data at the 2D class level or require a 3D model for particle discrimination.

### Denoising the particle images: cryoVAE

The first stage of our pipeline fits a VAE^19^ to the picked particles. We choose to use a standard beta-VAE as described here^20,21^. Β-VAEs are a type of generative model which aim to learn a latent, lower dimensional representation z of the input data x. The lower dimensional representation enables the VAE model to function as an effective de-noising strategy, faithfully reconstructing the underlying signal while ignoring the noise.

More formally, a β-VAE maximizes the Evidence Lower Bound (ELBO):

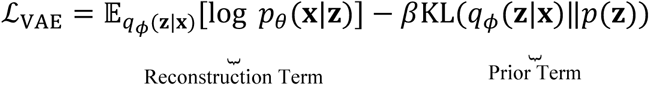

The ELBO consists of two terms: 1) the reconstruction term and 2) the prior term. The reconstruction term maximizes the likelihood of reconstructed data given the latent representation z and decoder model *p*_θ_(x | z). The prior term minimizes the difference between the encoded representation of the data, given by *q_φ_*(z | x), and a prior over the latent representation p(z), through minimizing the Kullback-Leibler (KL) divergence. Both the encoder *p*_θ_(x | z) and decoder *q_φ_*(z | x) are parameterized by neural network models. The hyperparameter β controls the relative strength of the prior regularization term compared to the reconstruction loss in the ELBO. Increasing β encourages the model to learn more disentangled, smoother latent representations at the cost of reconstruction fidelity.

We implement our β-VAE with a convolutional encoder and decoder. Further details can be found in the original implementation^19^. Following the standard VAE formulation, we choose to use an isotropic Gaussian prior p(z) =N (0, I). For reliable learning from the VAE and accurate reconstruction, we also noted that the input particle stacks should undergo additional low-pass filtering, down to 10Å. We fix the dimensionality of z (the latent space) to 20. We also chose to fix the value of β to 0.1 as we found this to provide a good balance between matching the prior and a sufficiently good reconstruction error for cryoEM datasets. We specifically do not optimize the choice of hyper-parameters (such as beta and size of latent dimensions) or architecture, as we ideally would like a solution that works without extensive hyper-parameter tuning for different datasets.

Our VAE model effectively acts as a denoising stage, producing reconstructed picked particle images that are denoised. We found that omitting this stage would cause the subsequent domain adaptation and particle classification stage to be less proficient due to high noise in the particle images. We stress that we do not use these reconstructed/denoised images for any data reconstruction. They are only used as an intermediate step to train our classifier, cryoDANN, to assign particle quality scores and selection. This avoids any potential artifacts in the final 3D maps that could arise from the smoothing of fine structural details caused by denoising.

### Calculating particle quality score: cryoDANN

In the second stage of the pipeline, we pass the denoised particles through an image classifier. This classifier (cryoDANN) is trained on the RELION class ranker training dataset (EMPIAR-10812), a large dataset of labelled class average images from a variety of different samples. This dataset consists of 18,051 2D class averages and we use the provided class scores (found in ‘features_normalised.star’) to annotate them as ‘good’ (rlnClassScore >0) and ‘bad’ (rlnClassScore <=0). This resulted in a set of 7213 good class averages and 10838 bad class averages. For each particle, the classifier outputs a score. This score can be used to discriminate a subset of good particles from the total set of picked particles (see Results). We use this labelled class ranker dataset as the source data, and the picked particles (to score) as the target dataset. To effectively increase the number of samples available in the target dataset we use data augmentation, including random horizontal and vertical flipping, and random 90° rotations. To further balance the two datasets, weighted random sampling was used to select samples from the source and target domain proportionally.

We use the same CNN classifier network as used by RELION class ranker^5^. However, naively applying a pre-trained classifier for particle filtering can be problematic if the particle images in the unseen dataset are far from the data distribution that the classifier was trained on. To mitigate this issue, we apply domain adaptation and fine tune the weights of our classifier model on denoised picked particles. The key to domain adaptation^22–24^ is to move the learned feature representation closer to the unknown true test distribution without degrading the performance of the model on the training data.

Formally, unsupervised domain adaptation (UDA) considers sets of input features X and target labels Y, drawn from a source domain D_S_ and a related target domain D_T_. We are given a set of labelled samples, where y_i_ is available from the source domain:

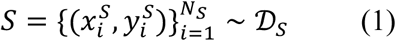

Likewise, we assume we have access to a set of unlabelled samples drawn from the target domain:

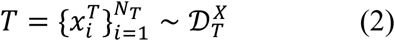

Ultimately, we would like to minimize the empirical risk R defined as:

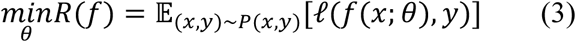

for the target domain T, where ℓ is a loss function (e.g. cross entropy) and *f* is a classifier with parameters θ, and 𝔼 is the expectation operator over all data samples. However, in UDA the target labels y are unknown and therefore this quantity cannot be directly minimized. Instead, we may use our knowledge from both the source and target domains to approximate this objective by combining 1) the empirical risk for the labelled source domain R_S_(f) and 2) simultaneously minimizing the discrepancy between the source and target distributions d(D_S_,D_T_):

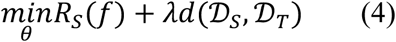

where λ is a hyperparameter.

In this work, we apply UDA to the problem of particle filtering. We treat the labelled data used for RELION class ranker as samples from the source distribution D_S_ and the new unlabelled particles as being sampled from the target distribution D_T_. We then train a new neural network f(·; θ) using UDA and minimizing the objective in equation 4.

For our choice of f (·; θ), we chose the domain adaptation neural network (DANN)^22^. DANN trains a neural network model with three components:

- Feature extractor h = g_h_(x; θ_h_) which maps the input data x to latent features h.
- Label classifier *ŷ* = *g_l_*(*h*; *θ_y_*) which maps latent features *h* to a label prediction *ŷ* for a given instance x_i_. In our specific case, this prediction is whether a given particle is good or bad.
- Domain classifier *d^* = *g_d_*(*h*; *θ_d_*) which maps latent features *h* to a domain prediction *d^*, which indicates whether a given sample xi came from the source or target domain.

DANN uses two loss components which correspond to the two terms in equation 4. To minimize the empirical risk, we compute the loss between the predicted label *ŷ* and the true source domain label y:

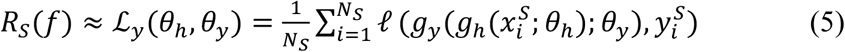

To align the feature distributions of the source and target domains, an adversarial objective L_d_ defined as:

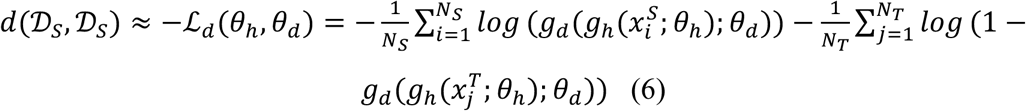

is used to encourage features h_i_ to be aligned and indistinguishable between source and target domains. The final objective to minimize is therefore:

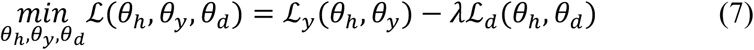

During training it is expected that domain loss will be high and domain accuracy low, while classification loss should be low and classification accuracy high. This would imply that the model can classify particle images well, while being invariant to domain-specific features.

### Pipeline for cryoEM data processing

We used a standardized processing pipeline on all datasets. This pipeline is a modified version of the relion_it.py script packaged with RELION^5^, rewritten using the CCP-EM Pipeliner (Iadanza *et al.*, under preparation, https://gitlab.com/ccpem/ccpem-pipeliner).

Automated processing at eBIC aims to provide live feedback as soon as data is acquired to monitor and inform adjustments to the acquisition process. For SPA experiments, this involves preprocessing stages which operate on individual micrographs, resulting in a set of picked particles. Preprocessing stages start with patch-based motion correction using MotionCor2^25^ (or RELION’s own implementation of motion correction, although MotionCor2 is used throughout here) with 5 patches in each dimension. CTF estimation is performed on each motion-corrected micrograph with CTFFind4^26^. Particle picking is performed with crYOLO^14^. The crYOLO general model trained on a diverse dataset is used, to remove the need for user intervention. crYOLO also estimates the size of each picked particle. The value of the 75th percentile of the first 10k picked particles is used to estimate the particle diameter. The box size used for particle extraction is taken to be 20% larger than the estimated particle diameter. Downscaling is then performed as described above, choosing box sizes from a list of sizes known to be performant in RELION^5^ classification. A mask diameter 10% larger than the estimated particle diameter is used for classification steps.

For efficiency of the pipeline, it is critical that the set of picked particles is filtered to remove contamination which may adversely affect 3D reconstruction. Currently, to perform this filtration, particles are subjected to reference-free 2D classification. To maintain the pipeline responsiveness, these 2D classifications must be performed on particle batches small enough that classification can complete in a reasonable amount of time. At eBIC, a particle batch size of 50k has been chosen. The particles are downscaled such that the Nyquist frequency post-binning is approximately 8.5 Å to further reduce the computational expense. Using 4 GPUs (NVIDIA V100) on a single node these jobs take approximately 20 mins to complete.

The subsequent particle sorting in our standard pipeline is performed at the level of 2D class averages, using the RELION ML-based class ranker tool termed ‘relion_class_ranker’. Particles estimated in the 2D classification step to have the highest likelihood of the pose of the selected 2D average are then used for 3D analysis steps. First, upon reaching threshold of 50,000 particles contributing to “good” 2D classes, an initial 3D model is generated^5^. Next, this model is used as reference for a 3D classification step where 200,000 “good” particles are selected.

In the eBIC SPA processing pipeline, further refinement is performed on particles assigned to the best 3D class with the best resolution estimate weighted by the cryoEF score^18^, if the resolution is better than 11 Å.

To test if cryoDANN can bring benefits to the SPA pipeline, we randomly selected 50 datasets from user visits in 2024 with reported 3D class resolution better than 13 Å from the standard eBIC pipeline. The rationale of this approach was to analyse both datasets of high quality and datasets close to the refinement threshold where relatively small resolution improvements may have an impact. The datasets were curated for 3D class quality to exclude those with unreliable 3D models leading to spurious resolution estimates. To test particle selection across these datasets, we implemented three test scenarios within the eBIC processing pipeline.

The first scenario is the pipeline with class ranker-based particle filtering (*f_a_*) discussed above.

In the second scenario, we replaced the class ranker particle selection step with a cryoVAE+cryoDANN particle selection. This step took each 50k particle batch, imposed orientation to each particle based on the pose estimate from the 2D classification job, ran cryoVAE on that batch to denoise the particle images, and then used cryoDANN to assign quality scores to each particle. The top half of the particles from the batch according to the cryoDANN score was selected. Re-batching, initial model generation and 3D classification was then performed as described above. This is filter *f_b_*.

The third scenario used a combination of both cryoVAE+cryoDANN and the RELION class ranker, ranking the particles in each batch by the product of the cryoVAE+cryoDANN score and the class ranker score, and then selecting the top 50% of particles. The rationale for this approach was that we would like to weigh down the contribution of the particles from the poor quality 2D classes, since the cryoVAE+cryoDANN approach alone does not discriminate between such images. This is filter *f_ab_*.

### Test datasets from EMPIAR

The method was tested on publicly available datasets downloaded from EMPIAR^4^. The dataset processing was done using the automated pipeline integrated at eBIC. Particle picking was performed using crYOLO^14^ and the particles split into batches of 50k to perform multiple downstream jobs in parallel. Our method was applied to these particle batches. Firstly, for each batch of particles (50k), the aligned stack of particle images (mrcs format) was saved using the in plane-transformations (rlnOriginX/Y and rlnAnglePsi in STAR file) by real space interpolation. This stack was denoised using the cryoVAE method. Following training, the output of cryoVAE comprises reconstructed (denoised) particle images, learned latent vectors, UMAP embedding of the latent vectors, a 2D histogram and a scatter plot of the UMAP embedding. The resulting denoised particle images were used as an input to cryoDANN to obtain the per particle quality score. Various thresholds for these scores were tested for their performance using the assessment metrics described below.

### Comparison with other methods and assessment metrics

At a selected threshold, cryoDANN selects the top N particles which can be used for further processing. We measured the performance of cryoDANN against sets of equal numbers of randomly selected particles and the top N particles suggested by class ranker^5^. The selected set of particles were used to perform 3D refinement. The metrics used to assess the performance were resolution (as defined by Rosenthal and Henderson, 2003^27^), B-factor (obtained by fitting a straight line through the Guinier plot) and angular distribution of the particles. The angular distribution of a given set of particles was assessed using the software cryoEF^18^, which calculates a coverage metric, referred to as the efficiency, that measures the quality of the angular distribution.

### Software availability

cryoVAE and cryoDANN plugins are included in the CCP-EM Doppio software (Figure 7). The software itself is separately available under an open-source MIT License from: https://gitlab.com/ccpem/cryovae https://gitlab.com/ccpem/cryodann

### Data availability

All the datasets which are used in this analysis are available from EMPIAR and the accession numbers are mentioned in the text. Release of the datasets used for testing the implementation of the method in eBIC live data processing pipelines is regulated by the Diamond Light Source user agreement: the data can be made publicly available after 3 years since data acquisition (year 2028 for the data used), providing no objections from respective data owners are received. An earlier release is possible upon request if data owners’ consent is received.

## Acknowledgements

We would like to thank Dr. Martyn Winn and Mr George Coldstream from Scientific Computing, STFC, for helpful discussions Dr. Satheesh Maheswaran for helpful discussions and management of this project during their time at Diamond. Agnel Praveen Joseph, Colin M. Palmer and Tom Burnley would like to thank the MRC for Partnership Grant MR/V000403/1, Sony Malhotra would like to acknowledge support from the Ada Lovelace Centre. Yuriy Chaban and Daniel Hatton would like to thank for support the EU/UKRI Fragment Screen Grant to Diamond, Agreement Number: 10059737. This work was funded by Diamond Light Source and Ada Lovelace Centre, UK Science and Technology Facilities Council. We acknowledge Diamond for access and support of the cryo-EM facilities at the UK national electron Bio-Imaging Centre (eBIC),

## Supplementary

Figure S1: Extracting the features from particle images. a. 2D histogram and b. scatter plot of the UMAP embedding of learned latent space vectors. The data shown is for a 50k particle subset for EMPIAR-10547, which were processed using the eBIC data processing pipeline.

Figure S2: Denoising of the particle images using cryoVAE. Before calculating the quality score per particle image, the particle stack is denoised using cryoVAE. The alignment of particle images is crucial to achieve meaningful training as seen in the quality of denoised images and the latent space visualisation.

Figure S3: The loss values and accuracies for the EMPIAR datasets obtained after cryoDANN training. For the four datasets the class and source domain loss and accuracies are plotted as the training progresses (increase in epoch number).

Figure S4: The fraction of particles in a given class above the cryoDANN score threshold, for manually selected ‘good’ classes and the remaining classes labelled as ‘others’ for the test datasets.

Figure S5: Pipeline performance comparison. a. Top: ratio of best 3D class resolution from the class ranker only particle filter *f_a_* to the combined filter *f_ab_* (blue) and the cryoVAE+cryoDANN only filter *f_b_*(orange). Larger values indicate better performance. Values above the dashed line at 1 indicate outperformance of the *f_a_* baseline, while values below indicate underperformance relative to the baseline. Bottom: ratio of cryoEF efficiency score for the best resolution 3D class produced by filters *f_b_* and *f_ab_* to that produced by *f_a_*. Again, larger values indicate better performance. *b and c.* The existing particle filter *f_a_* (blue, using the RELION class ranker) is compared to filter *f_b_*(orange, using cryoVAE and cryoDANN) and the combination filter *f_ab_* (green, using combination of *f_a_* and *f_b_*). The histogram and distribution (kernel density estimation) show the b. resolution and c. cryoEF score of the best 3D class for the 50 user datasets collected at eBIC.

Figure S6: Implementation of cryoVAE and cryoDANN in the CCP-EM software suite Doppio. a. The input form for the cryoVAE job where user can upload the particles and perform the denoising. b. cryoDANN input form which lets the users either use the raw particles or denoised particles obtained from cryoVAE jobs as an input to perform particle scoring. c. Output snapshots from cryoVAE and cryoDANN which show the images of low pass filtered particles, denoised particles, their latent space distribution and the distribution of particle scores.

Table S1: Particle filtering at different cryoDANN thresholds for EMPIAR-10024.

Table S2: Particle filtering at different cryoDANN thresholds for EMPIAR-10097.

Table S3: Particle filtering at different cryoDANN thresholds for EMPIAR-10093.

Table S4: Particle filtering at different cryoDANN thresholds for EMPIAR-11233.

Table S5: Particle filtering at different cryoDANN thresholds for EMPIAR-10255 (157k subset of particles).

Table S6: Particle filtering at different cryoDANN thresholds for EMPIAR-10255 (900k full set of particles).

Table S7: Automatic processing pipeline results for the three pipelines (a, b, and c) on the 50 eBIC datasets.

## References

1. Henderson, R. & Hasnain, S. ‘Cryo-EM’: electron cryomicroscopy, cryo electron microscopy or something else? IUCrJ 10, 519–520 (2023).

2. Henderson, R. From Electron Crystallography to Single Particle CryoEM (Nobel Lecture). Angewandte Chemie International Edition 57, 10804–10825 (2018).

3. Kühlbrandt, W. The Resolution Revolution. Science 343, 1443–1444 (2014).

4. Iudin, A. et al. EMPIAR: the Electron Microscopy Public Image Archive. Nucleic Acids Research 51, D1503–D1511 (2023).

5. Kimanius, D., Dong, L., Sharov, G., Nakane, T. & Scheres, S. H. W. New tools for automated cryo-EM single-particle analysis in RELION-4.0. Biochemical Journal 478, 4169–4185 (2021).

6. Punjani, A., Rubinstein, J. L., Fleet, D. J. & Brubaker, M. A. cryoSPARC: algorithms for rapid unsupervised cryo-EM structure determination. Nature Methods 14, 290–296 (2017).

7. Tang, G. et al. EMAN2: an extensible image processing suite for electron microscopy. J. Struct. Biol. 157, 38–46 (2007).

8. Moriya, T. et al. High-resolution Single Particle Analysis from Electron Cryo-microscopy Images Using SPHIRE. Journal of Visualized Experiments (JoVE) e55448 (2017) doi:10.3791/55448.

9. Sigworth, F. J. Principles of cryo-EM single-particle image processing. Microscopy (Oxf) 65, 57–67 (2016).

10. Lyumkis, D. Challenges and opportunities in cryo-EM single-particle analysis. The Journal of Biological Chemistry 294, 5181 (2019).

11. Unwin, P. N. T. & Henderson, R. Molecular structure determination by electron microscopy of unstained crystalline specimens. Journal of Molecular Biology 94, 425–440 (1975).

12. Drulyte, I. et al. Approaches to altering particle distributions in cryo-electron microscopy sample preparation. Acta Crystallogr D Struct Biol 74, 560–571 (2018).

13. Wagner, T. & Raunser, S. The evolution of SPHIRE-crYOLO particle picking and its application in automated cryo-EM processing workflows. Commun Biol 3, 1–5 (2020).

14. Wagner, T. et al. SPHIRE-crYOLO is a fast and accurate fully automated particle picker for cryo-EM. Commun Biol 2, 1–13 (2019).

15. J, Z., et al. A minority of final stacks yields superior amplitude in single-particle cryo-EM. Nature communications 14, (2023).

16. Clare, D. K. et al. Electron Bio-Imaging Centre (eBIC): the UK national research facility for biological electron microscopy. Acta Cryst D 73, 488–495 (2017).

17. McInnes, L., Healy, J. & Melville, J. UMAP: Uniform Manifold Approximation and Projection for Dimension Reduction. Preprint at 10.48550/arXiv.1802.03426 (2020).

18. Naydenova, K. & Russo, C. J. Measuring the effects of particle orientation to improve the efficiency of electron cryomicroscopy. Nat Commun 8, 629 (2017).

19. Kingma, D. P. & Welling, M. Auto-Encoding Variational Bayes. arXiv.org https://arxiv.org/abs/1312.6114v11 (2013).

20. Burgess, C. P., et al. Understanding disentangling in $\beta$-VAE. arXiv.org https://arxiv.org/abs/1312.6114v11 (2018).

21. Higgins, I. et al. beta-VAE: Learning Basic Visual Concepts with a Constrained Variational Framework. in (2017).

22. Ganin, Y. et al. Domain-Adversarial Training of Neural Networks. in Domain Adaptation in Computer Vision Applications (ed. Csurka, G.) 189–209 (Springer International Publishing, Cham, 2017). doi:10.1007/978-3-319-58347-1_10.

23. Farahani, A., Voghoei, S., Rasheed, K. & Arabnia, H. A Brief Review of Domain Adaptation. (2020). doi:10.48550/arXiv.2010.03978.

24. Kouw, W. M. & Loog, M. A Review of Domain Adaptation without Target Labels. IEEE Trans. Pattern Anal. Mach. Intell. 43, 766–785 (2021).

25. Zheng, S. Q. et al. MotionCor2: anisotropic correction of beam-induced motion for improved cryo-electron microscopy. Nat Methods 14, 331–332 (2017).

26. Rohou, A. & Grigorieff, N. CTFFIND4: Fast and accurate defocus estimation from electron micrographs. J Struct Biol 192, 216–221 (2015).

27. Rosenthal, P. B. & Henderson, R. Optimal determination of particle orientation, absolute hand, and contrast loss in single-particle electron cryomicroscopy. J Mol Biol 333, 721–745 (2003).

