## Supplementary material for "Automated filtering of particle images in single particle cryoEM"

[4]Research Complex at Harwell, Didcot, United Kingdom.

*co-corresponding authors

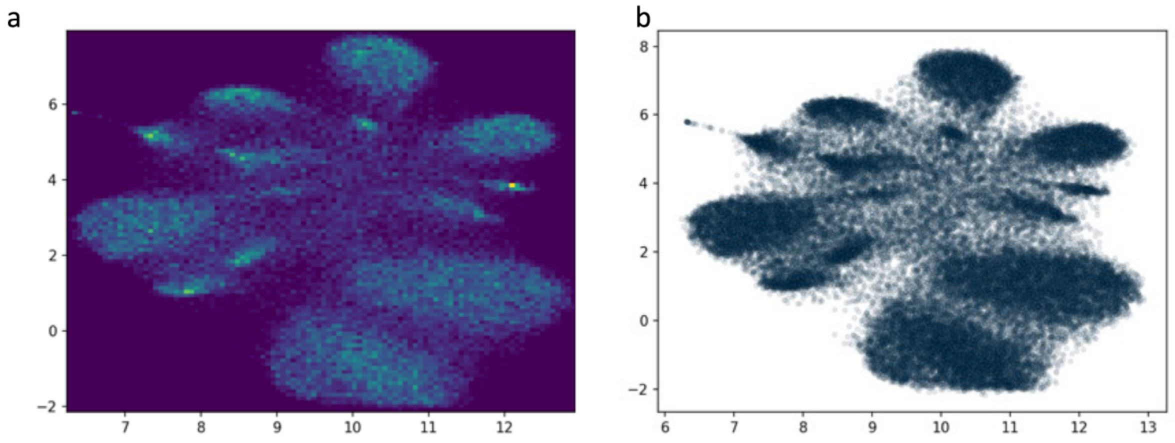


**Figure S1. Extracting the features from particle images. a.** 2D histogram and **b.** scatter plot of the UMAP embedding of learned latent space vectors. The data shown is for a 50k particle subset for EMPIAR-10547, which were processed using the eBIC data processing pipeline.

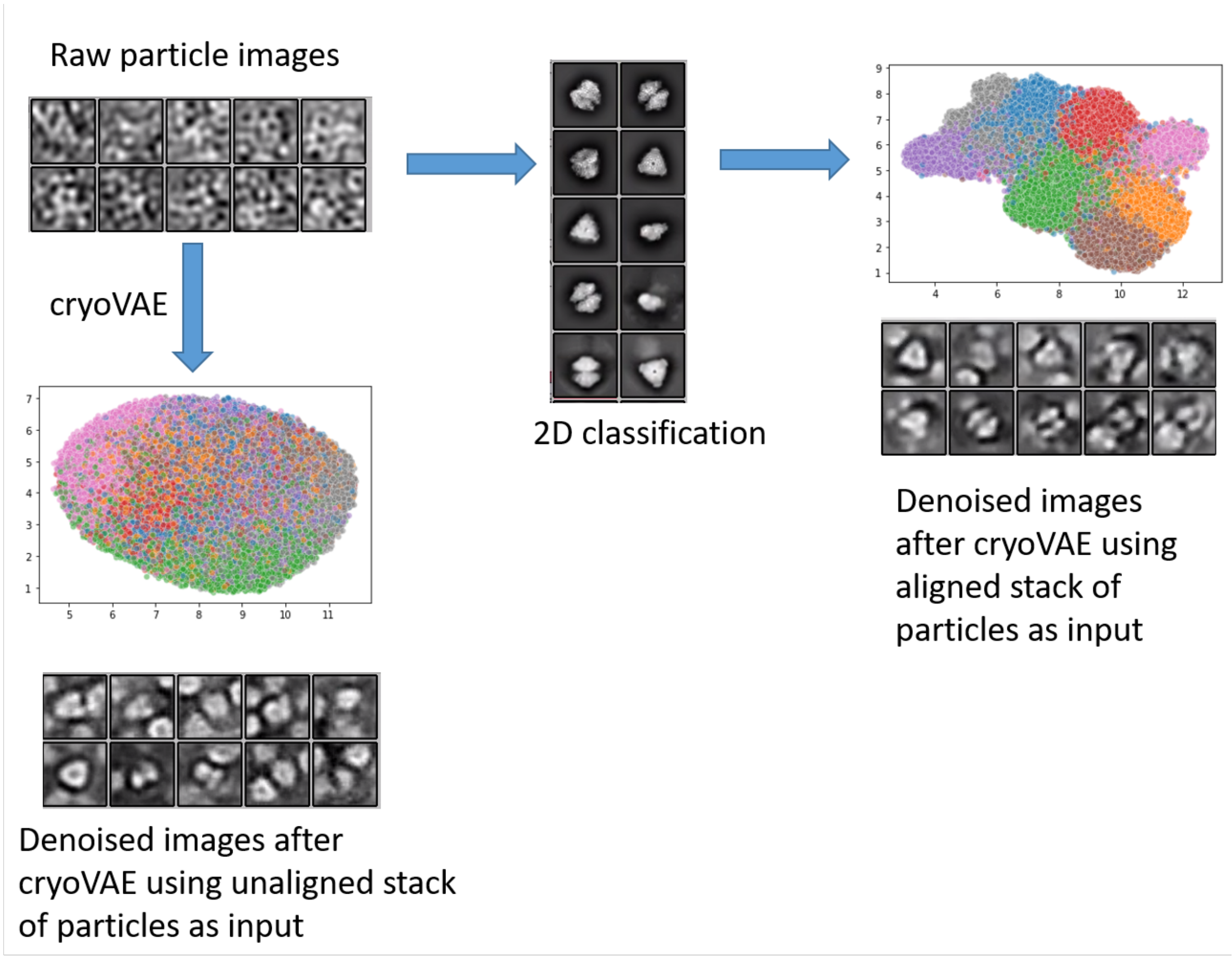


**Figure S2. Denoising of the particle images using cryoVAE.** Before calculating the quality score per particle image, the particle stack is denoised using cryoVAE. The alignment of particle images is crucial to achieve meaningful training as seen in the quality of denoised images and the latent space visualisation.

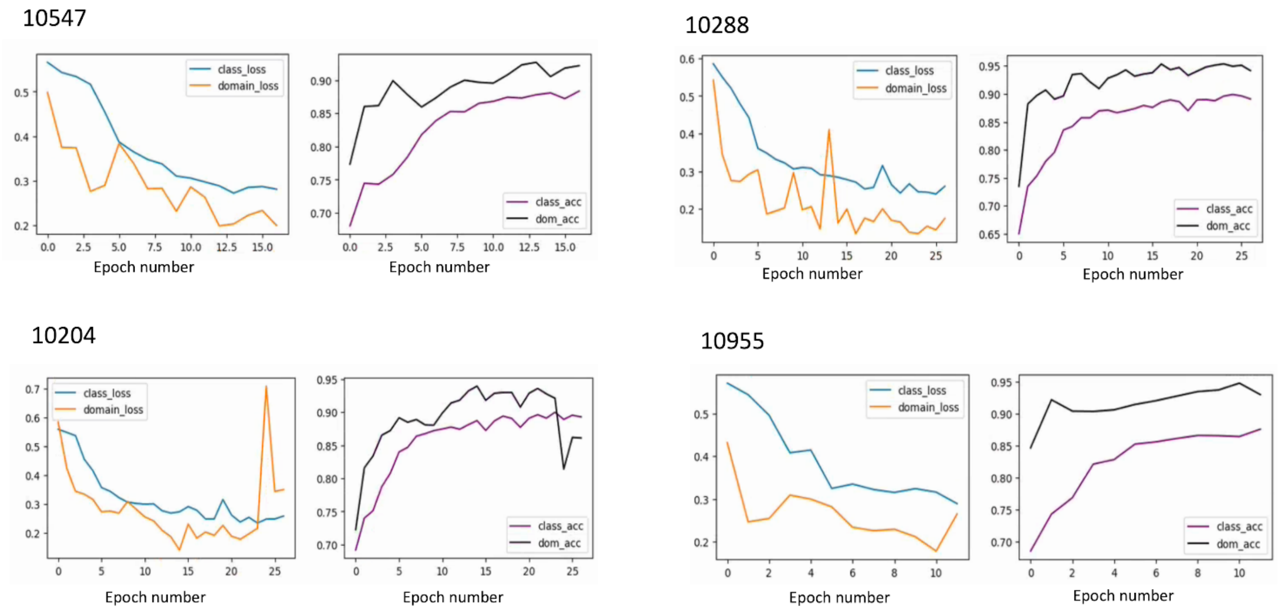


**Figure S3. The loss values and accuracies for the EMPIAR datasets obtained after cryoDANN training.** For the four datasets the class and source domain loss and accuracies are plotted as the training progresses (increase in epoch number).

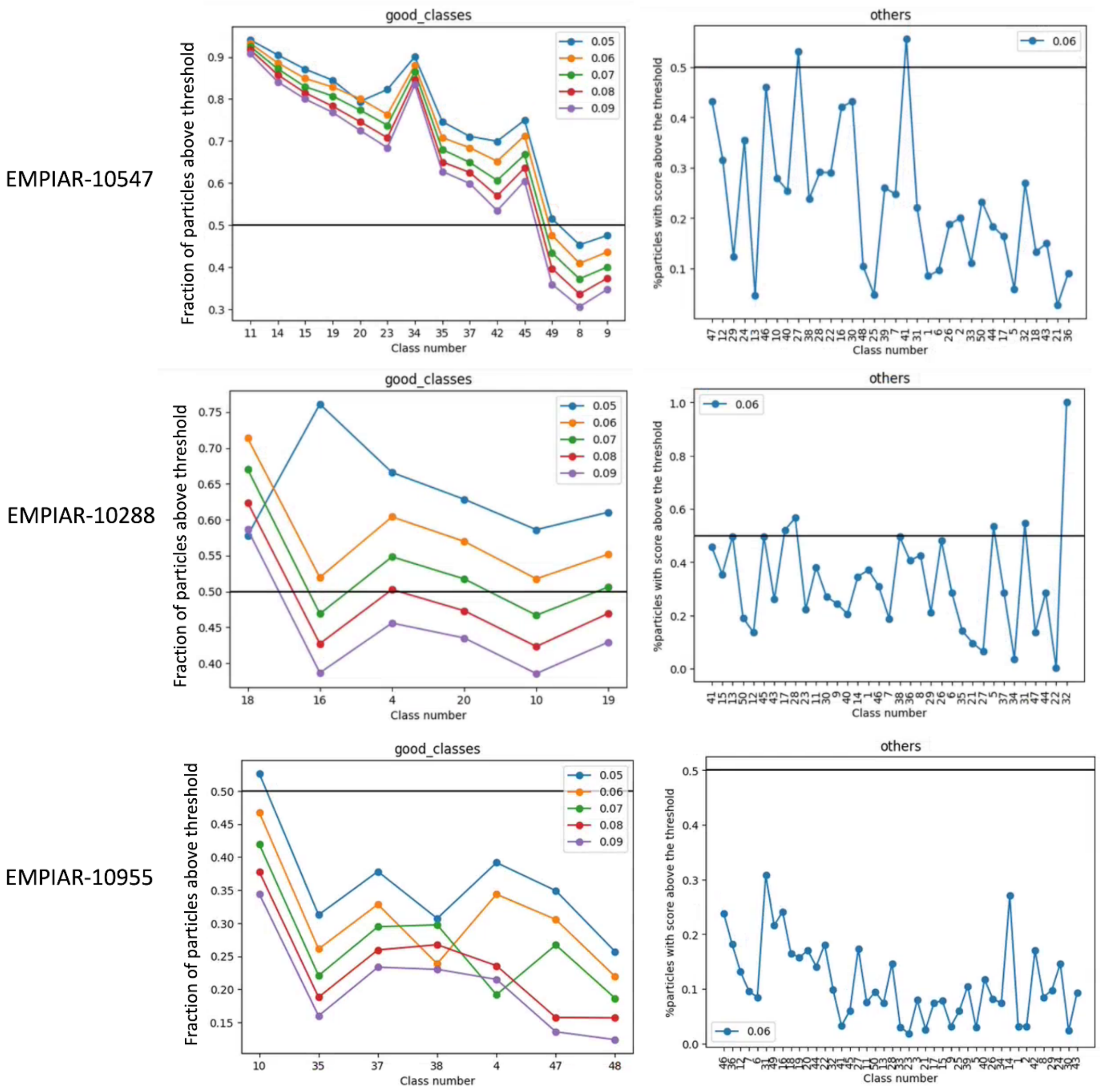


**Figure S4.** The fraction of particles in a given class above the cryoDANN score threshold, for manually selected ‘good’ classes and the remaining classes labelled as ‘others’ for the test datasets.

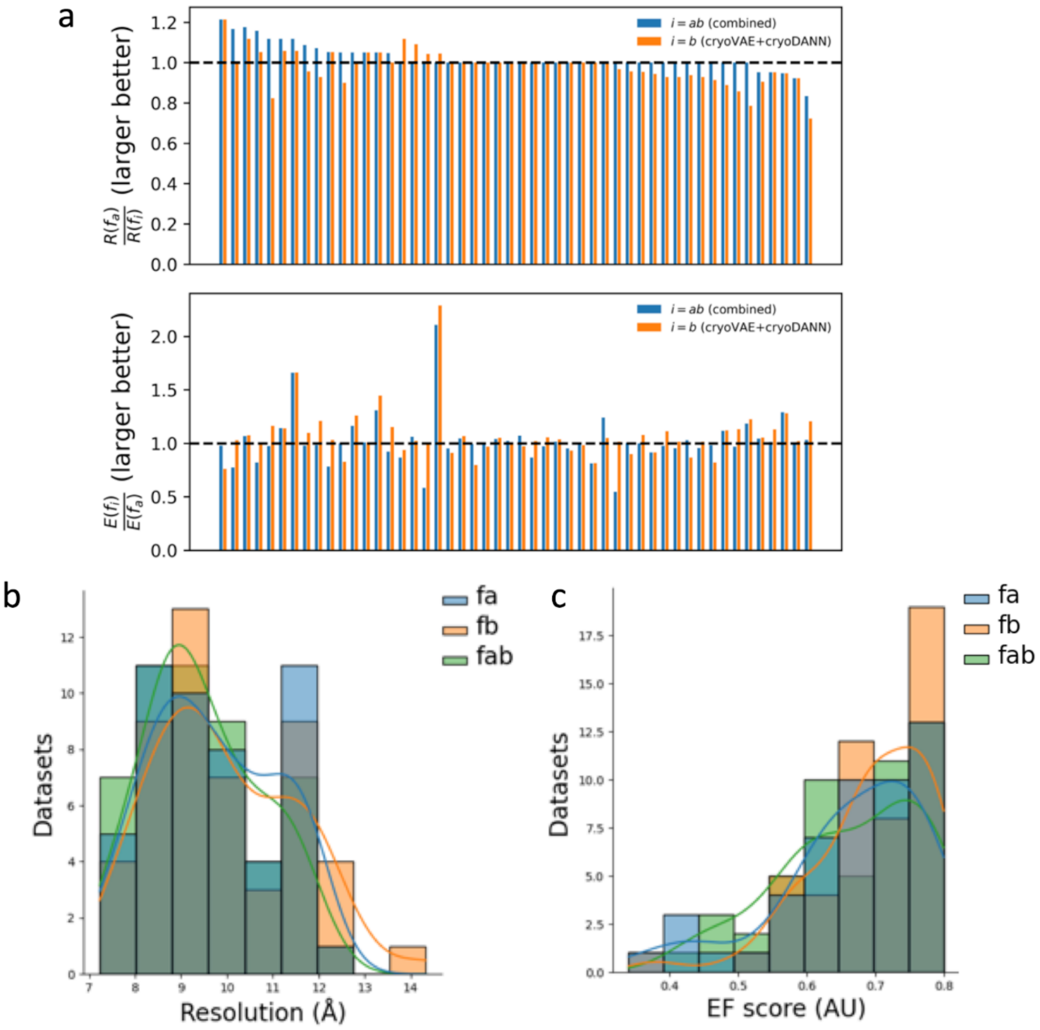


**Figure S5. Pipeline performance comparison. a.** Top: ratio of best 3D class resolution from the class ranker only particle filter $f_a$ to the combined filter $f_{ab}$ (blue) and the cryoVAE+cryoDANN only filter $f_b$(orange). Larger values indicate better performance. Values above the dashed line at 1 indicate outperformance of the $f_a$ baseline, while values below indicate underperformance relative to the baseline. Bottom: ratio of cryoEF efficiency score for the best resolution 3D class produced by filters $f_b$ and $f_{ab}$ to that produced by $f_a$. Again, larger values indicate better performance. **b.** and **c.** The existing particle filter $f_a$ (blue, using the RELION class ranker) is compared to filter $f_b$(orange, using cryoVAE and cryoDANN) and the combination filter $f_{ab}$ (green, using combination of $f_a$ and $f_b$). The histogram and distribution (kernel density estimation) show the **b.** resolution and **c.** cryoEF score of the best 3D class for the 50 user datasets collected at eBIC.

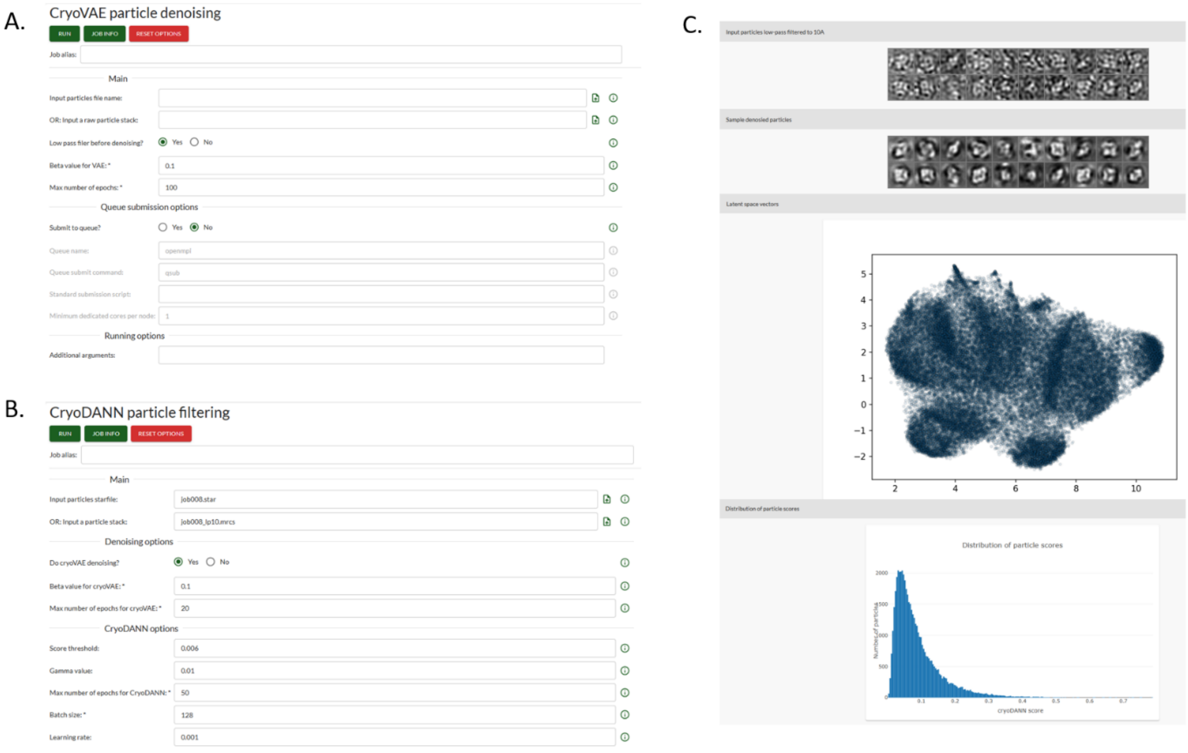

**Figure S6. Implementation of cryoVAE and cryoDANN in the CCP-EM software suite Doppio. a.** The input form for the cryoVAE job where user can upload the particles and perform the denoising. **b.** cryoDANN input form which lets the users either use the raw particles or denoised particles obtained from cryoVAE jobs as an input to perform particle scoring. c. Output snapshots from cryoVAE and cryoDANN which show the images of low pass filtered particles, denoised particles, their latent space distribution and the distribution of particle scores.

Table S1: Particle filtering at different cryoDANN thresholds for EMPIAR-10024.

| EMPIAR-10024 | | | | |
|---|---|---|---|---|
| **Set** | **Threshold** | **Number of particles** | **Resolution (Å)** | **(-)B-factor** |
| All particles | | 43585 | 4.24 | 191.92 |
| Top scoring | >=0.9 | 13535 | 4.73 | 132 |
| Top scoring | >=0.8 | 25278 | 4.39 | 178 |
| Top scoring | >=0.7 | 31462 | 4.29 | 215.74 |
| Top scoring | >=0.6 | 35705 | 4.29 | 195 |
| Random1 | | 13535 | 4.67 | 194 |
| Random2 | | 25278 | 4.39 | 180 |
| Random3 | | 31462 | 4.29 | 180 |
| Random4 | | 35705 | 4.29 | 178 |
| Low scoring | <=0.9 | 30050 | 4.29 | 168 |
| Low scoring | <=0.8 | 18307 | 4.67 | 184 |
| Low scoring | <=0.7 | 12123 | 5.44 | 176 |
| Low scoring | <=0.6 | 7880 | 7.15 | 505 |
| No denoising | 0.0003 | 26734 | 4.29 | 179 |
| No denoising | 0.0004 | 18602 | 4.39 | 183 |
| No denoising | 0.0005 | 12159 | 4.61 | 173 |
| Random no denoising | | 26734 | 4.39 | 190 |
| Random no denoising | | 18602 | 4.5 | 134 |
| Random no denoising | | 12159 | 4.7 | 118 |

Table S2: Particle filtering at different cryoDANN thresholds for EMPIAR-10097.

| EMPIAR-10097 | | | | |
|---|---|---|---|---|
| **Set** | **Threshold** | **Number of particles** | **Resolution (Å)** | **(-)B-factor** |
| All particles | | 130000 | 4.24 | 244.27 |
| Top scoring | >=0.9 | 31812 | 4.79 | 236 |
| Top scoring | >=0.8 | 66681 | 4.35 | 214 |
| Top scoring | >=0.7 | 88099 | 4.29 | 217 |
| Top scoring | >=0.6 | 101751 | 4.29 | 216 |
| Random | | 31812 | 4.79 | 220 |
| Random | | 66681 | 4.47 | 217.37 |
| Random | | 88099 | 4.35 | 212 |
| Random | | 101751 | 4.29 | 221.14 |
| Low scoring | <=0.6 | 28249 | 5.08 | 223.9 |
| Low scoring | <=0.7 | 41901 | 4.65 | 198.47 |
| Low scoring | <=0.8 | 63319 | 4.41 | 235.93 |
| No denoising | >=0.0009 | 78879 | 4.35 | 237.34 |
| No denoising | >=0.001 | 70351 | 4.24 | 214.32 |
| No denoising | >=0.0015 | 37645 | 4.72 | 207.55 |
| Random no denoising | | 78879 | 4.29 | 235.298 |
| Random no denoising | | 70351 | 4.35 | 226.87 |
| Random no denoising | | 37645 | 4.59 | 218.92 |

Table S3: Particle filtering at different cryoDANN thresholds for EMPIAR-10093.

| EMPIAR-10093 | | | | |
|---|---|---|---|---|
| **Set** | **Threshold** | **Number of particles** | **Resolution (Å)** | **(-)B-factor** |
| All particles | | 175344 | 3.74 | 140.11 |
| Top scoring | >=0.8 | 14762 | 5.5 | 155 |
| Top scoring | >=0.7 | 84725 | 4.08 | 151.56 |
| Top scoring | >=0.6 | 132775 | 3.85 | 149 |
| Top scoring | >=0.5 | 155390 | 3.76 | 139 |
| Random | | 14762 | 4.5 | 122 |
| Random | | 84725 | 3.8 | 142 |
| Random | | 132775 | 3.76 | 138 |
| Random | | 155390 | 3.76 | 141 |
| Low scoring | <=0.8 | 160552 | 4.26 | 255.88 |
| Low scoring | <=0.7 | 90589 | 4.38 | 230.19 |
| Low scoring | <=0.6 | 42539 | 4.86 | 239 |
| Low scoring | <=0.5 | 19924 | 6.31 | 439.73 |

Table S4: Particle filtering at different cryoDANN thresholds for EMPIAR-11233.

| EMPIAR-11233 | | | | |
|---|---|---|---|---|
| **Set** | **Threshold** | **Number of particles** | **Resolution (Å)** | **(-)B-factor** |
| All particles | | 42071 | 3.19 | 56 |
| Top scoring | >=0.95 | 17903 | 3.49 | 53 |
| Top scoring | >=0.9 | 28248 | 3.29 | 55 |
| Top scoring | >=0.85 | 32761 | 3.19 | 54 |
| Top scoring | >=0.8 | 35251 | 3.19 | 55 |
| Random | | 17903 | 3.56 | 67 |
| Random | | 28248 | 3.39 | 62 |
| Random | | 32761 | 3.29 | 56 |
| Random | | 35251 | 3.29 | 57 |

Table S5: Particle filtering at different cryoDANN thresholds for EMPIAR-10255 (157k subset of particles).

| EMPIAR-10255: 157k | | | | |
|---|---|---|---|---|
| **Set** | **Threshold** | **Number of particles** | **Resolution (Å)** | **(-)B-factor** |
| All particles | | 157224 | 3.7 | 222 |
| Top scoring | >=0.4 | 17803 | 3.9 | 160 |
| Top scoring | >=0.3 | 28216 | 3.84 | 167 |
| Top scoring | >=0.2 | 44682 | 3.77 | 214 |
| Top scoring | >=0.1 | 74383 | 3.71 | 159 |
| Random | | 17803 | 3.84 | 176 |
| Random | | 28216 | 3.84 | 163 |
| Random | | 44682 | 3.7 | 157 |
| Random | | 74383 | 3.7 | 177 |

Table S6: Particle filtering at different cryoDANN thresholds for EMPIAR-10255 (900k full set of particles).

| EMPIAR-10255: 900k | | | | |
|---|---|---|---|---|
| **Set** | **Threshold** | **Number of particles** | **Resolution (Å)** | **(-)B-factor** |
| Top scoring | >=0.95 | 53691 | 3.84 | 193 |
| Top scoring | >=0.94 | 66033 | 3.7 | 204 |
| Top scoring | >=0.93 | 78068 | 3.84 | 179 |
| Top scoring | >=0.9 | 112794 | 3.77 | 202 |
| Random | | 53691 | 4.05 | 220 |
| Random | | 66033 | 4.05 | 200 |
| Random | | 78068 | 4.05 | 220 |
| Random | | 112794 | 3.84 | 206 |

Table S7: Automatic processing pipeline results for the three pipelines (using filters $f_a$, $f_b$, and $f_{ab}$) on the 50 eBIC datasets discussed in Section 2.3. $R(f_i)$ indicates the lowest resolution of a 3D class produced with filter $f_i$. $E(f_i)$ indicates the cryoEF efficiency score of the same 3D class. The Nyquist resolution for the classes given the downscaling of the particles that occurs in the implementation of the pipelines is given for comparison.

| $R(f_b)$ **(Å)** | $R(f_a)$ **(Å)** | $R(f_{ab})$ **(Å)** | $E(f_b)$ | $E(f_a)$ | $E(f_{ab})$ | **Nyquist limit (Å)** |
|---|---|---|---|---|---|---|
| 9.87 | 11.99 | 9.87 | 0.57 | 0.75 | 0.74 | 6.99 |
| 9.16 | 10.24 | 8.70 | 0.78 | 0.72 | 0.77 | 7.25 |
| 8.98 | 10.03 | 10.03 | 0.69 | 0.74 | 0.64 | 7.11 |
| 10.26 | 11.19 | 11.19 | 0.75 | 0.74 | 0.78 | 5.13 |
| 10.04 | 10.63 | 9.51 | 0.79 | 0.69 | 0.79 | 7.53 |
| 9.82 | 10.40 | 9.30 | 0.71 | 0.42 | 0.71 | 7.36 |
| 10.29 | 10.83 | 10.29 | 0.68 | 0.66 | 0.52 | 6.43 |
| 9.49 | 9.99 | 8.63 | 0.70 | 0.71 | 0.58 | 7.91 |
| 8.76 | 9.20 | 8.76 | 0.78 | 0.78 | 0.78 | 7.67 |
| 7.79 | 8.18 | 7.79 | 0.63 | 0.44 | 0.57 | 6.82 |
| 8.04 | 8.39 | 8.39 | 0.78 | 0.78 | 0.46 | 8.04 |
| 7.59 | 7.93 | 7.93 | 0.79 | 0.34 | 0.72 | 7.27 |
| 11.80 | 11.80 | 10.11 | 0.79 | 0.77 | 0.60 | 5.90 |
| 11.48 | 11.48 | 11.48 | 0.70 | 0.76 | 0.73 | 6.22 |
| 8.35 | 8.35 | 7.97 | 0.69 | 0.60 | 0.55 | 7.31 |
| 7.73 | 7.73 | 7.73 | 0.56 | 0.53 | 0.55 | 6.76 |
| 9.11 | 9.11 | 9.11 | 0.60 | 0.75 | 0.75 | 8.26 |
| 11.46 | 11.46 | 11.46 | 0.77 | 0.80 | 0.79 | 4.77 |
| 9.54 | 9.54 | 9.54 | 0.78 | 0.74 | 0.77 | 3.98 |
| 11.94 | 11.94 | 11.94 | 0.69 | 0.69 | 0.70 | 5.47 |
| 11.49 | 11.49 | 11.49 | 0.72 | 0.74 | 0.79 | 5.27 |
| 10.85 | 10.85 | 10.85 | 0.59 | 0.58 | 0.50 | 5.43 |
| 9.57 | 9.57 | 9.11 | 0.50 | 0.40 | 0.46 | 7.97 |
| 8.54 | 8.54 | 8.54 | 0.67 | 0.63 | 0.61 | 6.76 |
| 8.74 | 8.74 | 8.74 | 0.76 | 0.73 | 0.74 | 4.37 |
| 11.29 | 11.29 | 11.29 | 0.69 | 0.74 | 0.71 | 8.47 |
| 9.05 | 9.05 | 9.05 | 0.78 | 0.80 | 0.79 | 7.16 |
| 8.79 | 8.79 | 8.79 | 0.61 | 0.75 | 0.61 | 5.13 |
| 8.86 | 8.86 | 8.86 | 0.66 | 0.63 | 0.78 | 6.65 |
| 8.59 | 8.31 | 8.31 | 0.76 | 0.75 | 0.41 | 8.05 |
| 7.55 | 7.23 | 7.23 | 0.72 | 0.80 | 0.80 | 6.93 |

| | | | | | | |
|---|---|---|---|---|---|---|
| 8.92 | 8.52 | 8.52 | 0.71 | 0.66 | 0.66 | 7.81 |
| 9.57 | 9.11 | 9.57 | 0.66 | 0.59 | 0.60 | 7.98 |
| 12.13 | 11.61 | 10.68 | 0.78 | 0.71 | 0.69 | 8.34 |
| 9.55 | 9.02 | 9.02 | 0.60 | 0.66 | 0.60 | 6.76 |
| 8.12 | 7.54 | 7.54 | 0.69 | 0.62 | 0.60 | 4.40 |
| 12.10 | 11.46 | 12.10 | 0.73 | 0.57 | 0.73 | 6.80 |
| 9.44 | 8.76 | 8.76 | 0.68 | 0.67 | 0.64 | 5.11 |
| 10.97 | 10.28 | 10.28 | 0.57 | 0.66 | 0.67 | 6.85 |
| 9.92 | 9.22 | 9.22 | 0.69 | 0.68 | 0.65 | 4.03 |
| 8.21 | 7.50 | 7.50 | 0.38 | 0.47 | 0.46 | 7.18 |
| 10.23 | 9.50 | 8.86 | 0.78 | 0.64 | 0.63 | 5.54 |
| 11.05 | 10.20 | 11.05 | 0.80 | 0.78 | 0.78 | 5.53 |
| 9.00 | 8.15 | 8.55 | 0.80 | 0.76 | 0.79 | 5.35 |
| 11.21 | 10.09 | 9.61 | 0.59 | 0.72 | 0.72 | 8.41 |
| 12.69 | 11.28 | 11.28 | 0.73 | 0.65 | 0.73 | 8.46 |
| 11.36 | 9.73 | 9.73 | 0.66 | 0.59 | 0.57 | 6.39 |
| 14.33 | 11.80 | 10.56 | 0.76 | 0.65 | 0.64 | 8.36 |
| 12.24 | 9.62 | 9.62 | 0.76 | 0.62 | 0.73 | 8.41 |
| 11.96 | 8.64 | 10.36 | 0.76 | 0.63 | 0.65 | 6.48 |
